# Nanoporous PEGDA ink for High-Resolution Additive Manufacturing of Scaffolds for Organ-on-a-Chip

**DOI:** 10.1101/2023.11.27.568937

**Authors:** Vahid Karamzadeh, Molly L. Shen, Houda Shafique, Felix Lussier, David Juncker

## Abstract

Polydimethylsiloxane (PDMS), commonly used in organ-on-a-chip (OoC) systems, faces limitations in replicating complex geometries, hindering its effectiveness in creating 3D OoC models. In contrast, poly(ethylene glycol)diacrylate (PEGDA-250), favored for its fabrication ease and resistance to small molecule absorption, is increasingly used for 3D printing microfluidic devices. However, applications in cell culture have been limited due to poor cell adhesion. Here, we introduce a nanoporous PEGDA ink (P-PEGDA) designed to enhance cell adhesion. P-PEGDA is formulated with a porogen, photopolymerized, followed by the porogen removal. Utilizing P-PEGDA, complex microstructures and membranes as thin as 27 µm were 3D-printed. Porogen concentrations from 10-30% were tested yielding constructs with increasing porosity and oxygen permeability surpassing PDMS, without compromising printing resolution. Tests across four cell lines showed >80% cell viability, with a notable 77-fold increase in MDA-MB-231 cell coverage on the porous scaffolds. Finally, we introduce an OoC model comprising a gyroid scaffold with a central opening filled with a cancer spheroid. This setup, after a 14-day co-culture, demonstrated significant endothelial sprouting and integration within the spheroid. The P-PEGDA formulation is suitable for high-resolution 3D printing of constructs for 3D cell culture and OoC owing to its printability, gas permeability, biocompatibility, and cell adhesion.

## Introduction

Organ-on-a-chip (OoC) systems have emerged as a promising alternative *in vitro* tool in various fields, including drug discovery, physiological monitoring, and personalized medicine^1–3^. These systems utilize microfluidics and tissue engineering to replicate the structure and function of human organs^4^. Leveraging 3D cell culture and microfluidics, OoC platforms have become an alternative to both *in vitro* 2D culture — due to their increased biological relevance — and *in vivo* animal testing, owing to their more controllable and reproducible nature. OoCs comprise microfluidic channels designed for 3D cell culture, with polydimethylsiloxane (PDMS) being most commonly used to fabricate OoCs^4,5^. However, it can be challenging to achieve complex structures with PDMS replica molding as fabrication is limited to replicating surface structures, while for example 3D printing allows for easier production of complex 3D structures with more design freedom^6^. Additional limitations of PDMS include the potential to absorb small hydrophobic molecules, which can result in inaccurate assessments of drug toxicity and efficacy^7^. Thus, researchers are exploring alternative natural and synthetic materials to enhance OoC capabilities. As the field of OoC technology rapidly evolves, 3D printing is emerging as a pivotal tool, offering new dimensions of flexibility and precision that surpass the capabilities of traditional PDMS replication methods^8–12^. This approach to fabrication not only circumvents the limitations inherent in PDMS-based designs but also paves the way for the creation of more complex and functionally diverse OoC systems. In addition, compared to classical microfabrication methods, the cost per device does not increase with increasing complexity or batch size, making it easier to customize and rapidly iterate to improve the design. Moreover, the technical knowledge required to operate 3D printers can be acquired quickly and easily compared to most other manufacturing methods due to the largely automated nature of the process^13^. However, layer-by-layer 3D printing methods face a trade-off between build height and print time, and based on the pixel size of common digital light processing (DLP) 3D printers (2-40 µm), cannot produce large polymer structures with geometrical features at the sub-micrometer scale, such as micro- and nanoporous scaffolds. Sub-micrometer scale 3D printing has been reported by using methods such as direct laser writing by 2-photon photopolymerization, but scalability remains a challenge due to low production rates, one-point-at-a-time photopolymerization scanning speeds, and highly specialized instrumentation^14,15^.

Recent studies on 3D printing have explored nanoporous materials formed by salt porogen leaching^16,17^, aqueous two-phase emulsion^18^, and polymerization-induced phase separation (PIPS)^19,20^. Each approach relies on different mechanisms for pore formation, producing different types of pores with varying size distribution and that include interconnected pores, blind pores, and closed pores, in different ratios. If interconnected pores predominate, then it is characterized as open porosity, and if blind and closed pores predominate, it is closed porosity. The salt porogen leaching method is limited to creating micropore sizes due to the salt particle size, while two-phase emulsion requires careful optimization of surfactants to ensure the immiscible phases remain uniform throughout the printing process, which may not be compatible for cell studies. PIPS in particular is well suited for photopolymerization 3D printing, as it utilizes a stable miscible porogen in the ink and can create complex nanoporous structures with tunable porosity. However, most of these techniques necessitate post-processing with expensive instruments, such as supercritical drying^19,20^. For example, Dong *et al*. demonstrated a method that combines vat-photopolymerization (VP) 3D printing of 2-hydroxyethyl methacrylate (HEMA) monomer and PIPS to create 3D objects with nanoporosity by removing cyclohexanol and 1-decanol porogens^20^; note that low molecular weight porogens may also be a solvent, and vice-versa. These 3D-printed nanoporous structures showed improved cell attachment, and presented excellent compatibility of heterogeneous material VP as the precursor phase is miscible while the photopolymerized phase is immiscible, causing a phase separation of the solid cured ink from its uncured counterpart, making PIPS an exceptional method of forming nanostructures via VP. However, the use of short chain HEMA created brittle structures that depend on supercritical drying for preservation, making the process impractical for many applications. Moreover, a high concentration of a photoinitiator (PI) (4% Irgacure 819) known for its toxicity and potentially long printing times for PIPS constitute further downsides.

Poly(ethylene glycol) (PEG) is a biocompatible and hydrophilic biomaterial that finds wide use in tissue engineering and pharmaceutical applications^21^. By incorporating acrylate or methacrylate functional groups, PEG becomes photocurable and suitable for VP 3D printing. PEG diacrylate (PEGDA), with molecular weight (MW) of 250-6000 Da, has become widely used in 3D printing as a biocompatible 3D printing material^22,23^. PEGDA with molecular weight of 250 Da can be 3D-printed without dilution and forms a water-impermeable, transparent polymer that can be chemically functionalized. Formulations with higher polymer weight of ∼500-6000 Da must be mixed with water for 3D printing and form hydrogels with excellent biocompatibility properties. Compared to PDMS, PEGDA-250 is easier and less expensive to manufacture and exhibits innate resistance to absorption of small molecules, and suitable for drug testing applications^13^. However, low-MW PEGDA suffers from low cell attachment compared to PDMS^22^.

Nonetheless, several groups have developed PEGDA-250 formulations for high-resolution 3D printing, and for cell culture applications notably using phenylbis(2,4,6-trimethylbenzoyl) phosphineoxide (BAPO) as the PI^23–27^. Folch and colleagues demonstrated the ability to 3D-print microchannels with a cross-section of 27-µm-wide and 1-mm-tall using low-MW PEGDA with isopropylthioxanthone (ITX) as the photoabsorber (PA). Nordin and colleagues printed microchannels with an even smaller cross-section of 18 × 20 μm^2^ using 2-nitrophenyl phenyl sulfide (NPS) PA^24,27^. However, prior to their use in cell culture, these formulations required minimum 24 h washing and surface plasma treatment to make them biocompatible and allow for cell attachment^22,24^. The International Organization for Standardization (ISO) standards for cell biocompatibility have been formulated, but only one study adopted them when studying the cytotoxicity of the material for only one cell line^22^. Therefore, there is an unmet need for a biocompatible, easy-to-manufacture PEGDA ink that promotes cell attachment.

High resolution nanoporous ink development thus relies on a balance between short monomers and porogens that are be easily removed. Although low-MW PEGDA has largely been favored for high-resolution VP, PEGDA-250 and PEGDA-258 are insoluble in water^28^, thus limiting miscible porogen selection. Commonly used solvents like methanol can evaporate during 3D printing, while larger porogens reduce print layer adhesion as the monomer size is relatively small. An emerging alternative with high PEGDA-compatibility is the use of PEG as a porogen^29^, which is miscible and non-photocurable due to lacking terminal acrylate groups. PEG can be synthesized at varying MWs, making it suitable for tunable pore sizes, and for cell-based applications.

Here, we present a novel biocompatible porous PEGDA (P-PEGDA) ink that is easy to 3D-print, and which includes an inert PEG porogen easily removed by washing, resulting in nanoporous prints. We demonstrate P-PEGDA inks with different degrees of porosity, characterize 3D printability, gas permeability, and diffusion of fluorescent dyes into the prints. We further present a comprehensive cytotoxicity study on four cell lines with varying susceptibility according to ISO standards and confirm the beneficial effects of nanoporosity on cell attachment. Next, we introduce a novel OoC system directly made by 3D-printing featuring a central chamber for receiving a cancer spheroid, a gyroid scaffold for culturing cells, and a separation barrier with microchannels and bi-directional capillary stop valves on multiple levels to permit selective seeding of spheroid and cells surrounding the spheroid in all directions. We report long-term co-culture of spheroids and cells, observe cell proliferation, bi-directional 3D cell migration, and endothelial sprouting within the spheroid.

## Results

### Characterization and Design Criteria for P-PEGDA Ink

Current inks for VP 3D printing mainly rely on a monomer and a PI, which results in solid, nanoporous structures. In this study, we present a novel formulation that includes a porogen compatible with hydrophilic PEGDA monomer and PI. By simply washing the 3D-printed object with water, the porogen can be easily removed, leaving behind nanoporous structures **(Figure 1a)**. Our approach is faster, more cost-effective, and simpler compared to previously reported methods that depend on supercritical drying to controllably remove the porogen^20^. This approach enables the printing of high-resolution microstructures with nanoporosity, making it suitable for OoC device fabrication.

**Figure 1.**
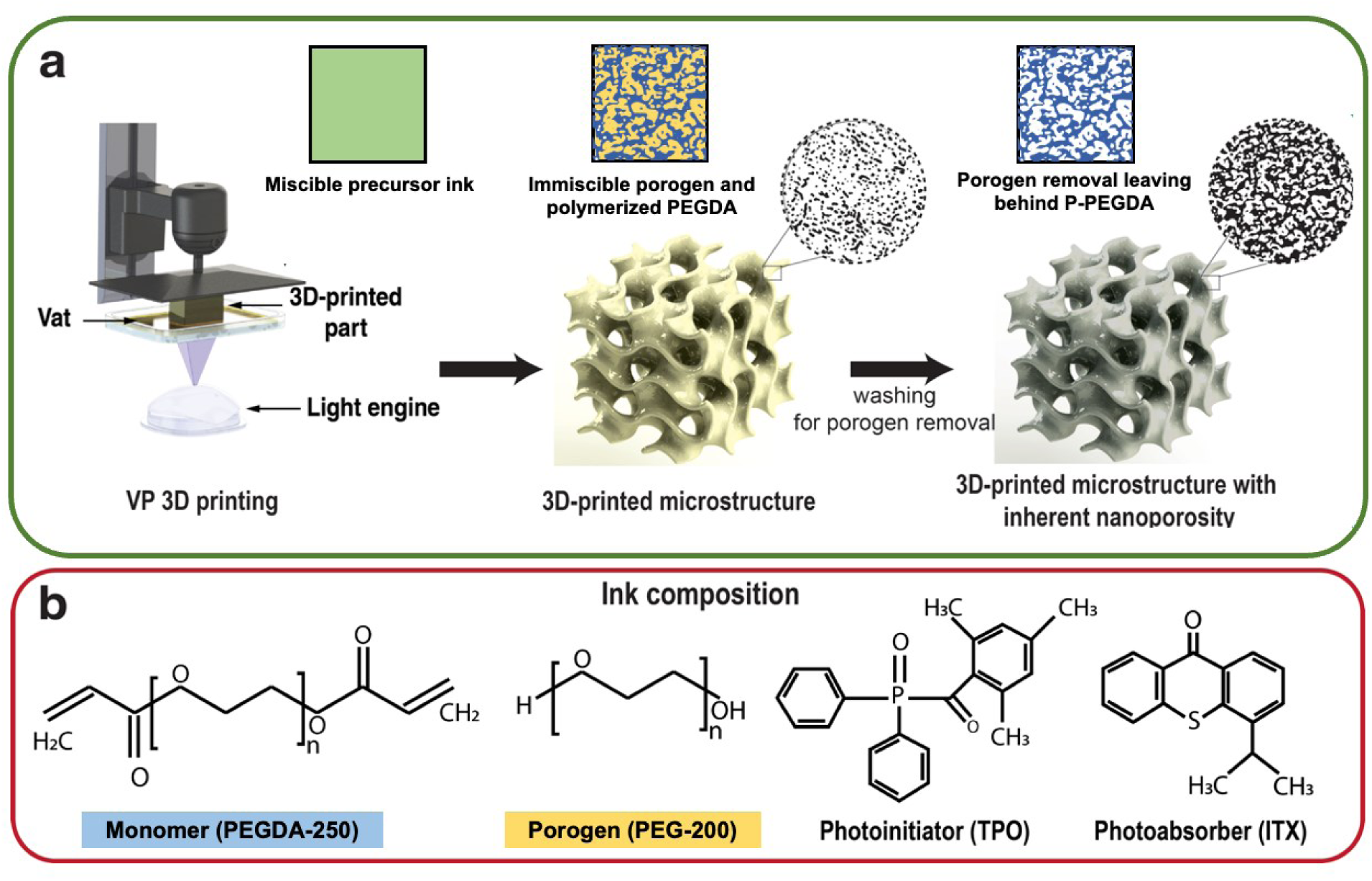
Schematic of the 3D printing process and the ink composition. **(a)** The fabrication process starts with 3D printing of an object to achieve microporosity, followed by washing steps to leach out the PEG porogen to form nanopores. The black areas in the images indicate the porosity of the structures. **(b)** P-PEGDA ink composition including PEGDA-250 monomer, PEG-200 porogen, TPO PI, and ITX PA.

### The PEGDA-based formulation suitable for OoC applications had to meet the following criteria

(a) 3D printable with high resolution, (b) having high transparency for high-resolution fluorescence microscopy, (c) gas permeable for closed channel devices, (d) being biocompatible and suitable for long-term cell culture, and (e) having sufficient cell attachment. Based on these criteria, we developed an ink consisting of PEGDA-250 as the hydrophilic monomer, PEG-200 as the porogen, TPO as the PI, and ITX as the PA **(Figure 1b)**. These criteria are essential to ensure that the ink could be used effectively for the fabrication of complex OoC devices.

We included PEG-200 as a porogen in our formulation for its ability to dissolve in hydrophilic monomers and diffuse out of the cured structure, leaving behind pores. It is also non-toxic, FDA-approved, and has been widely used in biomedical applications^30,31^. The rationale behind using a porogen is to create a porous structure which can allow for oxygen exchange and facilitate the removal of toxic and unreacted components in 3D-printed structures. In our case, PEG-200 was found to be compatible with the PEGDA-250 and the PI, making it a suitable choice for making nanoporous inks and improving cell attachment.

The choice of PI for our PEGDA-based formulation for OoC applications requires careful consideration of several factors, including adequate absorbance at 385 nm, low cytotoxicity, and compatibility with the monomer. After narrowing the options to lithium phenyl-2,4,6-trimethyl benzoyl phosphinate (LAP), BAPO, and diphenyl(2,4,6-trimethylbenzoyl)phosphine oxide (TPO), we chose TPO as the optimal PI for several reasons. Firstly, TPO is highly soluble in PEGDA-250 and has shown low cytotoxicity compared to BAPO^32,33^, meeting the requirements of our formulation. Although LAP has the lowest cytotoxicity, it has very low solubility in PEGDA-250 (**Figure S1, Supporting Information**). Secondly, TPO has a maximum absorption at 380 nm, but with a wavelength window extending to up to 425 nm making it suitable for VP 3D printing, which generally operates with a 365-405 nm UV light source. Lastly, we observed that 3D-printed samples using TPO as the PI exhibited less yellowing compared to samples 3D-printed with BAPO, resulting in 3D-printed structures more suitable for a wider excitation range in fluorescence microscopy (**Figure S2, Supporting Information**). Overall, the use of TPO as the PI in our ink formulation satisfies the necessary criteria for our OoC application and provides optimal performance.

To achieve high-resolution 3D printing with the P-PEGDA ink, a PA was necessary to improve the vertical resolution and prevent clogging of conduits caused by optical penetration in the preceding layers. ITX was chosen as the PA due to its high efficiency at 385 nm, which is the optimal wavelength for our printing process. Additionally, ITX has been previously used with PEGDA-250 for high-resolution 3D printing and has low cytotoxicity, making it suitable for OoC applications^24^. In addition, compared to other PAs such as NPS, ITX is less cytotoxic and yellowish (**Figure S2, Supporting Information**). Therefore, the use of ITX in our ink formulation helps to meet the necessary criteria for our OoC application, including 3D printability with high resolution, high transparency for fluorescence microscopy, and cytocompatibility. The efficiency of the photoabsorber and ink is notably high at a wavelength of 385 nm, harmonizing with the UV spectrum of 385 nm DLP light engines (**Figure S3a, Supporting Information**). It’s notable that changes in the concentrations of PEG-porogen do not impact the absorbance of the ink (**Figure S3b, Supporting Information**).

In VP 3D printing, the resolution of embedded features in the Z direction is mainly dictated by the depth of light penetration into the ink. Characterization of the light penetration depth of the ink is crucial to optimize the printing parameters such as layer exposure time and layer thickness. We measured the penetration depth of light for different porogen concentrations at various exposure times to optimize the printing parameters for high accuracy in the Z direction. As shown in **Figure 2a**, the addition of PEG porogen slightly increased the cure depth; meanwhile, the penetration depth (given by the slope of the line) does not change, which is in concordance with the UV-Vis spectra of inks with different PEG concentrations. Interestingly, the minimum energy to cure is decreased with more porogen; the porogen dilutes the monomer, so this could be attributed to the higher PI to monomer ratio, leading to faster curing. Alternatively, the PEG porogen may act as a plasticizer, which can increase the mobility of the polymer chains in the material, leading to a more open and less dense network structure, which may also contribute to the increase in the penetration depth of light.

**Figure 2.**
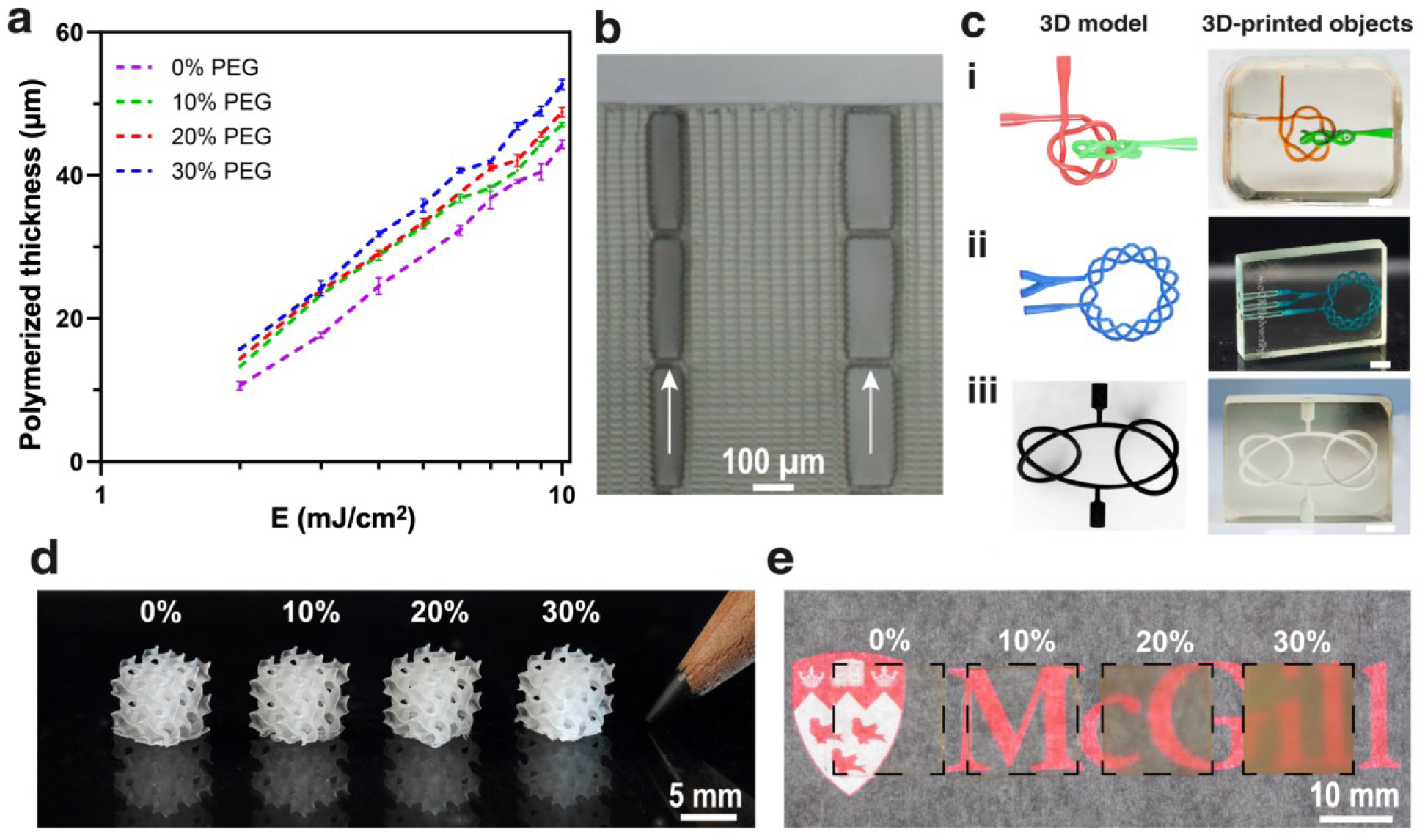
3D printability of P-PEGDA ink. **(a)** Polymerized thickness of P-PEGDA with different concentrations of porogen. Representative of three independent experiments (n=3). The error bar represents the standard deviation. **(b)** 3D printed stacked microchannels separated with 27-µm-thick membranes (indicated by white arrow). **(c)** 3D printed complex and intricate models with 10% porogen: **(i)** two-(2,5)-torus knots passing each other with a diameter of 340 µm, **(ii)** (3,13)-torus knot with a diameter of 240 µm, **(iii)** grammy knot with a diameter of 300 µm. Scale bar: 2 mm. **(d)** 3D-printed gyroid with a dimension of 5 mm^3^ at different porogen concentrations. **(e)** 3D printed 3D squares at different porogen concentrations; higher porogen concentrations result in decreased transparency of the 3D-printed parts.

Another contributor to high resolution printability is the low viscosity of the ink formulations, which allows for fast refilling of the vat between print layers and reduces the suction force between the 3D printed part and the flexible vat bottom, thus favoring 3D printing of delicate features. The P-PEGDA formulation is based on PEGDA-250 monomer which has an inherently low viscosity (∼16 mPa s) supplemented with PEG-200 porogen which has a higher viscosity (∼55 mPa s), but comparatively low when in a mixture with the monomer. Indeed, while higher porogen concentration slightly increased the viscosity of the formulation (**Figure S4, Supporting Information**), it still remains much lower than many alternative inks for biomedical applications. The ability to 3D print monolithic and embedded microchannels is crucial for microfluidic and OoC applications. These microchannels serve as the fundamental building blocks for complex OoC devices. To demonstrate the printability of P-PEGDA’s low viscosity and low penetration depth, we characterized the 3D printing resolution by 3D printing a stack of channels separated with a thin 27 µm membrane, using a formulation containing 10% porogen **(Figure 2b)**. The successful printing of these intricate structures, membranes and microfluidic channels shows the potential of the P-PEGDA ink for high-resolution 3D printing of complex microstructures for applications such as OoC devices.

The high-resolution printing capability of our ink enabled us to achieve excellent printability and accuracy, producing complex and precise geometries such as torus knots that would be difficult to achieve with other fabrication methods. Torus knots are constructed mathematically by taking a circle in the torus and moving it around the torus in a certain way. The resulting knot is defined by two integers (p,q), which determine its overall geometry. The values of p and q indicate the number of times the knot winds around the minor and major axis of the torus, respectively. Using P-PEGDA (10%), we were able to successfully 3D print a grammy knot with a diameter of 300 µm, a (3,13)-torus knot with a diameter of 240 µm (**Video S1, Supporting Information**), and two-(2,5)-torus knots passing each other with a diameter of 340 µm **(Figure 2c)**. Moreover, we demonstrated the versatility of our ink by 3D printing objects like triply periodic minimal surfaces such as gyroids (**Figure 2d**) and 3Dbenchy (**Figure S5, Supporting Information**), using P-PEGDA with different porogen concentrations. The transparency of P-PEGDA ink is maintained for concentrations of 0% and 10% porogen; however, it decreases for concentrations above 10% (**Figure 2e**). This reduction in transparency can be attributed to the increased presence of pores within the material, which results from higher porogen concentrations. A higher presence of disordered pores leads to light scattering and increased opacity of the 3D printed material^20,34,35^. These findings suggest that a 10% porogen concentration is optimized for applications requiring both optical transparency for fluorescence microscopy and porosity.

### Effect of porogen concentration on material properties

To visualize the pores formed in the 3D-printed object, the printed samples were immersed in water to leach out the porogen and create voids within the structure. We performed a quantitative analysis of the pore size distribution using image analysis to evaluate the pore size dependency in the 3D printed parts to porogen concentration. We varied the concentration of PEG porogen in the ink to evaluate its effect on pore size distribution. We measured 40 individual pores from scanning electron microscope (SEM) images and analyzed the pore size distribution. We found that the pore size distribution was narrow for lower porogen concentrations but became broader at higher concentrations which might be related to the increasing diffusion of porogen during the printing process **(Figure 3a-b)**. Additionally, the average pore size increased from 5 nm to 30 nm for a porogen concentration of 10-30% compared to the 0% PEG samples. This observation is consistent with previous studies that have shown that higher porogen concentrations can lead to larger pore sizes in 3D-printed structures^18^. As the PEG concentration increases, the porogen-to-monomer ratio also increases, resulting in larger pore sizes. To determine if these pores impacted the surface roughness of the 3D printed P-PEGDA formulations, we mapped the surface topography with varied concentrations of porogen using atomic force microscopy (AFM). As expected, the AFM surface plots showed that higher porogen concentration led to greater areal surface roughness, with 30% and 0% porogen featuring a nanoroughness of 44.1 ± 3.8 nm and 15.2 ± 2.6 nm, respectively, **Figure 3c, Figure S6, Supporting Information**.

**Figure 3.**
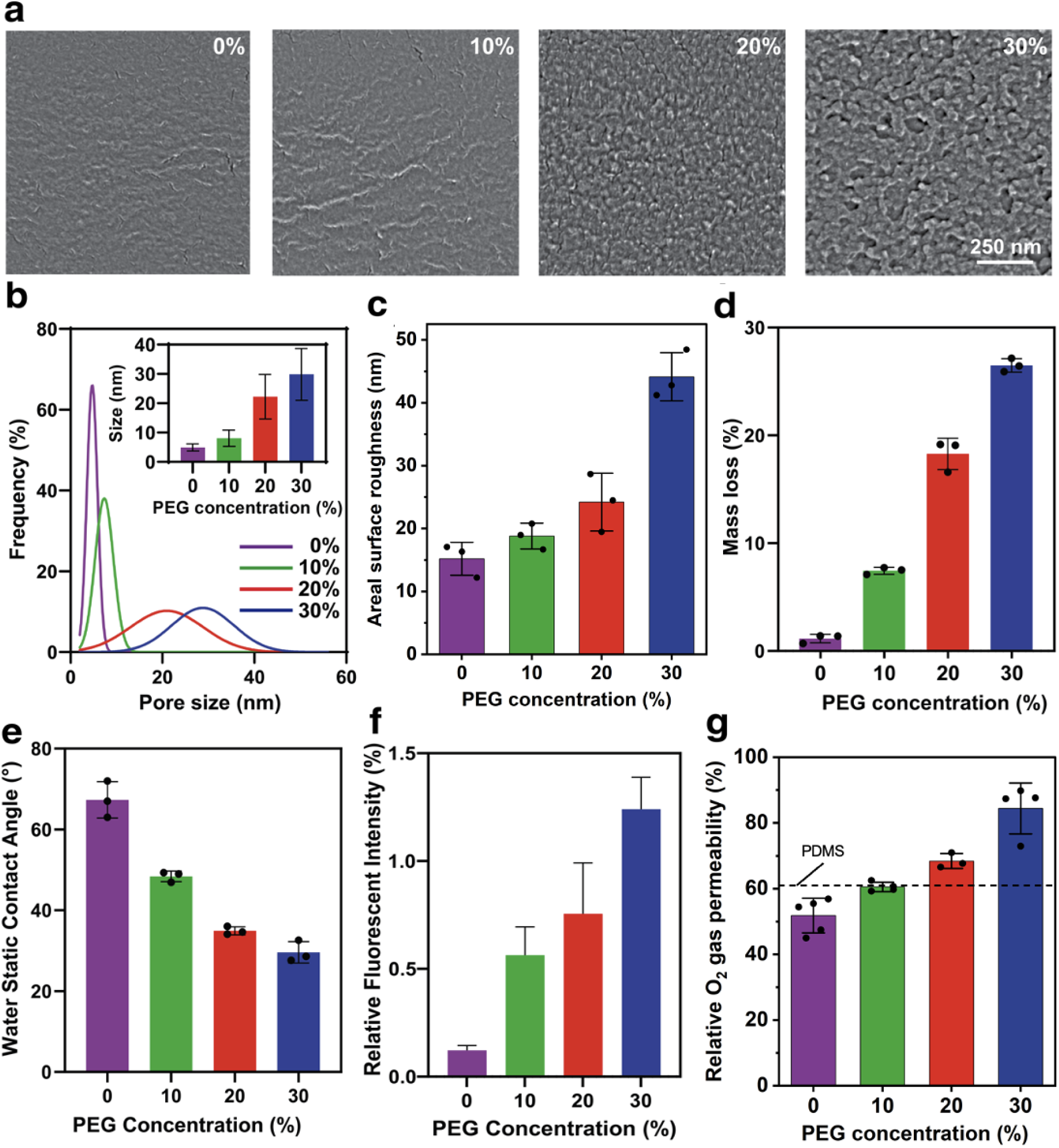
Characterization of P-PEGDA ink for different concentrations of PEG porogen. **(a)** SEM images show the porosity at different porogen concentrations. **(b)** Pore size distribution in the 3D-printed parts with P-PEGDA at different concentrations of porogen. Insert shows the average size for different concentrations. In each case, 40 random pores were analyzed. Mean of n=40 measurements and error bars are standard deviations. **(c)** Areal surface roughness of 3D printed P-PEGDA at different concentrations of porogen from AFM topographical profiles. **(d)** Mass loss of P-PEGDA after 48 h of incubation in DI water. Mean of n=3 measurements and error bars are standard deviations. **(e)** Static water contact angle for P-PEGDA for different porogen concentrations. Mean of n=3 measurements and error bars are standard deviations. **(f)** Relative fluorescent intensity after 6 min for different porogen concentrations. **(g)** Relative oxygen gas permeability of 3D-printed P-PEGDA barrier caps (used to seal a tube) with different porogen concentrations, compared to a tube without barrier (see Figure S8 in Supporting Information for further details). The dashed line shows the permeability of a PDMS cap. Mean ± standard error is presented.

To investigate the effect of porogen concentration on weight loss in our formulations, we fabricated 3D-printed disks with varying porogen concentrations and measured their mass immediately after printing. Following crosslinking, the disks were submerged in water for 48 h to ensure complete removal of the porogens, and then dried and massed again to determine their net mass loss. Our findings indicate that mass loss increased as porogen concentration increased, with disks containing 30% porogen experiencing a mass loss percentage of 26%, while those with 0% porogen only lost 2% of their initial mass **(Figure 3d)**. This trend is due to the greater amount of porogen available for removal during incubation in water in disks with higher porogen concentrations.

By incorporating a porogen into PEGDA, which is inherently hydrophilic before crosslinking, and subsequently removing it through washing after polymerization, a more hydrophilic material with a reduced contact angle is obtained (**Figure 3e**). The porogen’s addition creates pores within the material, increasing its surface area and disrupting the matrix structure, potentially providing additional sites for water molecule interaction. This process may lead to a rise in hydrophilic sites, resulting in enhanced hydrophilicity of the material. Furthermore, the porogen’s low molecular weight could contribute to the disruption of the PEGDA matrix, yielding a more disordered and less dense network. This factor may also play a role in increasing the material’s hydrophilicity and reducing its contact angle. The contact angle decreased from 67° contact angle for the 0% porogen objects to 29° contact angle for the objects 3D printed with 30% porogen. In our recent work^36^, we showed that a hydrophilic ink (CCInk) can be produced by adding acrylic acid to the formulation, reducing the contact angle to 35°. The P-PEGDA ink could potentially also be used for the fabrication of capillary microfluidic devices, as an increase in surface roughness caused by surface nanoporosity increases the wettable area and subsequently decreases the water static contact angle, likely due to capillarity and hemiwicking^37,38^.

In addition to the SEM analysis, we also carried out a diffusion test utilizing 3D-printed rods to investigate the diffusion of small molecules through the P-PEGDA structure. We washed the samples for 24 h to remove the porogen. Samples were then immersed in a solution of 10 μM fluorescein in MilliQ water, and fluorescence images were acquired by confocal microscopy near the middle height of the polymeric rod. By probing a single plane far from both the top and bottom parts of the pillar, negligible diffusion from fluorescein molecules is expected from either the top or bottom of the printed rod within the probed time. Hence, the increase in fluorescence intensity is solely associated with the radial diffusion of fluorescein molecules toward the center of the printed rod. Here, we measured the fluorescence intensity inside the rod by generating a ring-shaped mask. The mask was separated into distinct sectors (n=50) to assess the homogeneity in diffusion, and hence porosity, while omitting the center of the rod (**Figure S7, Supporting Information**). In addition, a ring-shaped mask was generated to assess the intensity of the bulk solution and used as normalizing factor to account for photobleaching during imaging. While we observe an increase in fluorescein diffusivity with the addition of porogen, the diffusion rate in 3D-printed P-PEGDA structures even with 30% porogen remains low and reaches only ∼1.2% of the solution concentration. (**Figure 3f**). This suggests that fluorescein does not diffuse through the PEGDA matrix (as expected), and that the pores might not be interconnected. However, the significant mass loss observed indicates the successful removal of most of the porogen, possibly due to the smaller molecular size of PEG200 compared to fluorescein, and possible swelling by ethanol. The mass loss suggests permeability for smaller molecules such as oxygen. To assess oxygen permeability, we placed cellulose test strips dip-coated with resazurin in gas impermeable tubes capped with 3D printed P-PEGDA discs. The reduction of resazurin can be visualized as the strips change color from blue to pink in the absence of oxygen. The tubes were placed under a vacuum for 48 hours, and gas permeable caps allowed for oxygen to exit and the strip to change color, while impermeable caps maintained the blue strip color. The oxygen permeability assay provided insights into the gas permeation properties of P-PEGDA ink. Specifically, inks containing porogen concentrations higher than 10% demonstrated higher permeability (>68%) compared to 1:10 PDMS (>61%), which is well-known to be gas permeable, and interestingly, the 10% porogen formulation exhibited an oxygen permeability close to that of PDMS, **Figure 3g, Figure S8, Supporting Information**. Altogether, the 10% porogen concentration balances oxygen permeability and optical transparency, rendering it an attractive choice for organ-on-a-chip systems.

Considering our findings, it’s important to acknowledge that while SEM imaging and mass loss data provide valuable insights into the porosity of our P-PEGDA structures, they offer a limited view of the pore interconnectivity. To gain a comprehensive understanding of porosity and connectivity, further studies are necessary to establish the relationship between porogen concentration on one hand, and the molecular weight, composition, and diffusion rate of various molecules on the other hand. Porosity and permeability are important properties that will guide the use of P-PEGDA for various biomedical applications.

### Cytocompatibility

To further demonstrate the advantages of our P-PEGDA ink, we evaluated its cytotoxicity based on two ISO standards, namely ISO 10993-12:2009 involving media transfer and ISO 10993-5:2009 involving co-culture with cells^39^. The former standard involves incubating cell media with the material for an extended period, retrieving the media, and exposing the media to cultured cells to measure the potential cytotoxicity of soluble elements released by the material. The latter is the direct co-culture of cells in the presence of the material immersed in the same cell culture media to measure the potential cytotoxicity of both released soluble elements as well as direct contact with the material. ISO 10993-12:2009 provides guidance on the determination of the potential for a medical device to cause skin irritation, while ISO 10993-5:2009 provides guidance on the determination of the potential for medical devices such as implants to cause cytotoxicity. The ISO 10993-12:2009 standard typically recommends a surface area to volume ratio of 6 cm²/mL for solid devices, a guideline that we adhered to in conducting our extraction assay. We selected several cell types ranging widely in their sensitivity, including immortalized human fibroblast cell line IMR-90, breast cancer cell line MDA-MB-231, immortalized human kidney cell line 293T, and human umbilical vein endothelial cell line HUVEC, for a cytotoxicity assay. The PrestoBlue assay was used to evaluate cell viability for its compatibility with time-course measurement in live cells.

We tested the biocompatibility of the P-PEGDA by measuring cell viability for the above-mentioned four cell lines using 3D printed disks that were washed for 24 h with 70% ethanol following their manufacturing. The samples likely contain unreacted components such as irradiated PIs and PAs that are removed during the post-manufacturing washing to render them biocompatible. Our results showed cell viability exceeding 80% for all cell lines for both ISO 10993-12:2009 and ISO 10993-5:2009 standard experiments, which meets the ISO standard test criteria of a viability >70% (**Figure 4a-b, Figure S9, Supporting Information)**. For example, the cell viability of HUVEC after 24 h was 90% for the ISO 10993-5:2009 standard. Moreover, we found that the cell viability was generally higher for the ISO 10993-12:2009 than the ISO 10993-5:2009 as expected. It is important to note that the post-manufacturing washing step is crucial in removing unreacted components that may be present in the 3D-printed part and that could diffuse, be taken up by cells, and affect their viability. The optimal washing time may vary depending on the application; for instance, a microfluidic device designed for a rapid cell assay may require little to no washing, while long-term culture of sensitive cell types such as induced pluripotent stem cells (iPSC) may require long washing times. Overall, our findings of high biocompatibility confirm the suitability of P-PEGDA for OoC applications.

**Figure 4.**
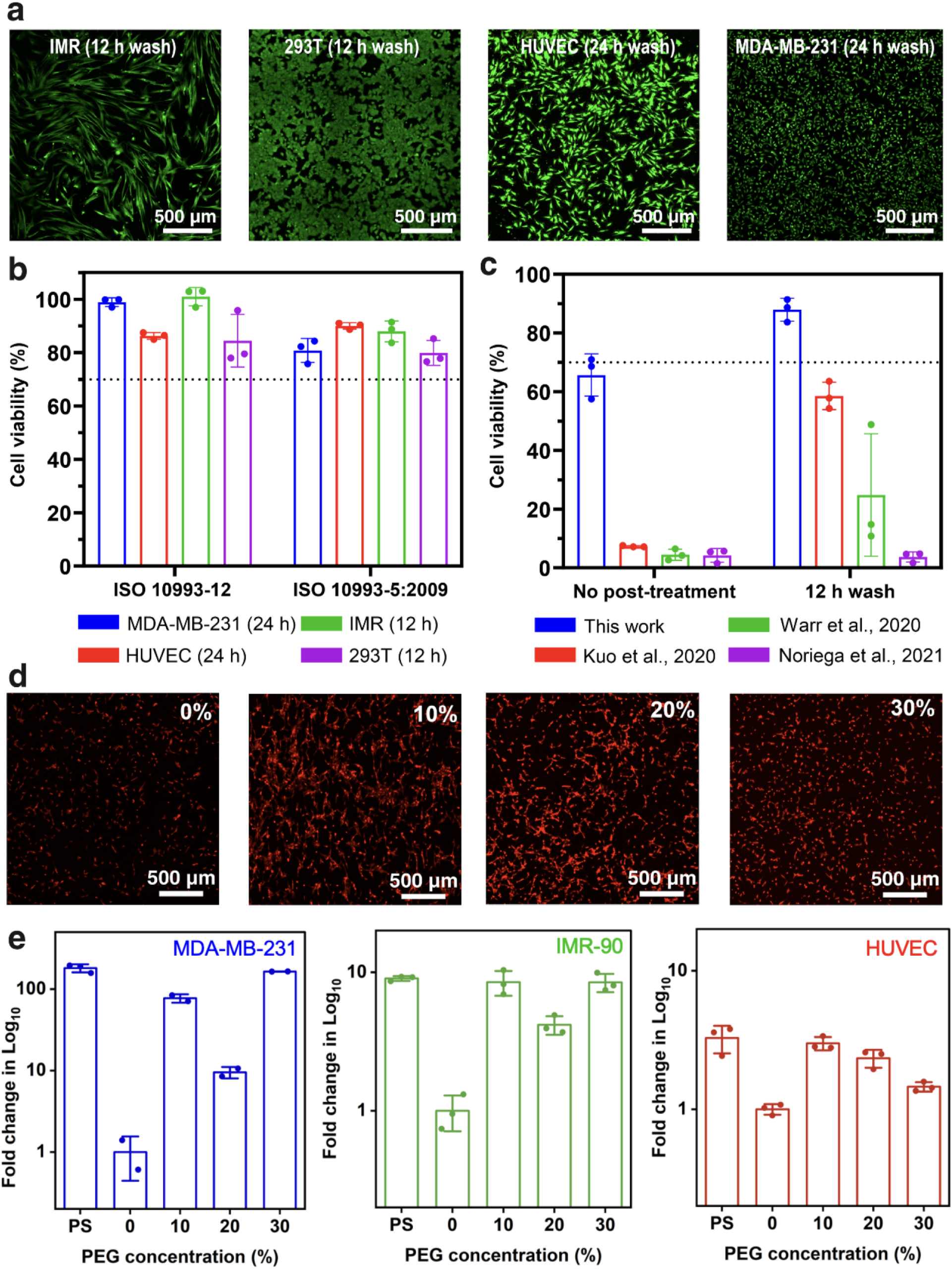
Cytotoxicity of, and cell attachment on, P-PEGDA. **(a)** Fluorescence images of 4 cell lines (IMR-90, 293T, HUVEC, and MDA-MB-231) co-cultured with P-PEGDA and imaged by fluorescence microscopy following 48 h of culture in a well-plate in the presence of 3D-printed samples washed in 70% ethanol for 12 h or 24 h. **(b)** Cell viability for various cell types for 3D-printed disks with different washing times, assessed by incubation with cell media (ISO 10993-12) and direct co-culture with the cells (ISO 10993-5). The line in **(b)** and (**c**) represents the minimum viability in accordance with the standards. **(c)** A comparison of the cytotoxicity between P-PEGDA ink and previously published low-MW PEGDA-based inks. Cell viability for IMR-90 cells after 48 h of culture in a well-plate along with 3D-printed samples without and with prior incubation in 70% EtOH for 12 h. Cell viability is highest for P-PEGDA both without and with washing compared to prior PEGDA-250 formulations by Kuo et al.^24^ (containing 0.4 % BAPO PI and 0.8 % ITX PA), Noriega et al.^27^ (containing 1% BAPO PI and 2% NPS PA), and Warr et al.^22^ (containing 1% BAPO and 0.38 % Avobenzone). **(d)** Cell attachment for HUVEC cells cultured on 3D-printed wells with P-PEGDA at different porogen concentrations. **(e)** Cell coverage per projected area on plasma-treated polystyrene (PS) (tissue culture plastic) and 3D-printed wells with different porogen concentrations for MDA-MB-231, IMR-90 and HUVEC cell lines. The error bar represents standard deviation of n=3 independent experiments.

Next, we investigated the impact of washing time and the ITX PA on the biocompatibility of 3D-printed objects for 293T, as it scored the lowest cell viability among the 4 cell lines tested (i.e., ∼80%). Unreacted components during the printing process may affect the biocompatibility of the printed object, and our results showed improved cell viability following the washing of the print with 70% ethanol. Cell viability increased from less than 70% without washing to higher than 80% after 12 h of washing (**Figure S9, Supporting Information**). The addition of 0.8% ITX did not negatively affect cell viability, as the samples with and without ITX showed similar increases in cell viability after washing. These results suggest that a 12 h washing period is sufficient to ensure biocompatibility of 3D-printed objects.

We compared the cytotoxicity of our P-PEGDA ink with previously published low-MW PEGDA-based inks using the direct co-culture method with IMR-90 cells (ISO 10993-5) (**Figure 4c** and **Figure S10, Supporting Information**)^22,24^. Notably, the P-PEGDA demonstrated superior biocompatibility when compared with the other low molecular weight PEGDA-based formulations even without any washing steps. In addition, the P-PEGDA formulation consistently exhibited a higher cell viability (**Figure S10, Supporting Information**). This advantageous performance can be attributed to the use of TPO PI, which here replaces the commonly used BAPO PI that is known for its cytotoxicity. Next, we cultured cells within 3D-printed enclosed channels (**Figure S11, Supporting Information**), and imaged them by fluorescence microscopy, further lending support to the biocompatibility of P-PEGDA and underlining its transparency and low autofluorescence. PEGDA ink has garnered significant attention in biomicrofluidics and OoC applications due to its 3D printability and biocompatibility. However, existing PEGDA inks lack both cell attachment and nanoporosity. Thus, prior research enhanced cell attachment on nonporous 3D printed PEGDA samples by plasma treatment, which necessitates the utilization of costly instrumentation^22^. Additionally, this approach may prove ineffective for internal structures that remain unexposed to plasma. In this study, we find that nanoporosity resulting from a PEG porogen promotes cell attachment, consistent with another study on phase-change-induced nanoporosity that observed similar effects^20^.

To assess cell attachment, we seeded 3D-printed P-PEGDA microwells with fluorescently tagged MDA-MB-231, IMR-90 and HUVEC cells. Cell attachment within each microwell was visualized using fluorescence microscopy after 24 h, and cell coverage was quantified via image analysis. For all cell types, a greater area of cell coverage was observed for P-PEGDA formulations with porogen compared to those without porogen, which suggests that the presence of pores indeed influences cell adhesion to the 3D-printed microwells, and in some cases, the adhesion was similar to that observed for plasma-treated polystyrene (PS) (tissue culture plastic), the gold standard material for 2D cell culture, **Figure 4d-e**. Notably, a 77-fold increase in cell attachment was observed in P-PEGDA (10%) for MDA-MB-231 microwells compared to the 0% PEG condition.

Although the trend of surface adhesion varies for different cell lines, the tested cell lines consistently show greater adhesion for the porous scaffold (10-30% PEG) compared to the 0% PEG counterparts. Consequently, 3D scaffolds with nanoporosity demonstrate promising potential for OoC applications involving 3D cell culture. The increased cell adhesion on nanoporous scaffolds can be attributed to several factors. First, the nanoporous scaffolds provide a larger surface area for cells to adhere^40^. Second, the nanoporous topography can enhance cell anchorage, by allowing more filopodia to attach firmly to the surface^41^. These findings have also been observed in HepG2 cells^20^, indicating suitability for a range of cell types. Interestingly, the microwells with the highest attachment of HUVECs also visually demonstrated cell alignment, which suggests that the nanoporosity led to cellular organization. To further investigate the alignment, we converted the fluorescent microscopy images into binarized masks and measured Feret’s angle of cells with respect to an arbitrary axis. The 10% PEG formulation of P-PEGDA exhibited a narrow directionality, while HUVECs on other formulations and on plasma-treated polystyrene (PS) had a broadly distributed cell alignment, which suggests that P-PEGDA 10% could play a role in cell organization based on nanoporosity, and outperforms PS in this regard, **Figure S14, Supporting Information**.

### Organ-on-a-chip with P-PEGDA

Following our findings that P-PEGDA with 10% porogen promoted better cell adhesion while preserving transparency, we sought to evaluate its potential for OoC application with a tumor-on-a-chip model. A common OoC design is based on culturing spheroids (or organoids) surrounded by stromal and endothelial cells that induce vascularization^42–45^. The initial segregation and compartmentalization of spheroids and endothelial cells relies on capillary stop valves (CSVs) also called phaseguides, which are realized using parallel microchannels on either side of the spheroid. Typically spheroids are seeded in the center with a hydrogel precursor that remains confined by the valves, followed by filling of the channels with endothelial cells that can then proliferate from the bottom into the spheroid and vascularize it via angiogenesis^46^. Yet cells *in vivo* within their native environment in tissues and organs are typically surrounded by stromal and endothelial cells from all sides.

Here, we developed a novel OoC system that provides connectivity on all sides comprising (i) a gyroid serving as a 3D scaffold for culturing stromal or endothelial cells with (ii) a vertical hole in the center forming a chamber for receiving a spheroid, (iii) a circular separation wall between the chamber and the gyroid, and (iv) 30 microchannels (250 µm in diameter) across the wall on 3 levels radiating in 360 degrees and terminated in both central and radial directions by CSVs, **Figure 5a-c**. Gyroids are particularly suitable for tissue engineering and regenerative medicine, owing to their porous and interconnected shape and high surface-to-volume ratio^47,48^. The interconnectivity of the gyroid facilitates cell migration and tissue integration which is advantageous for tissue engineering^49,50^. The dual CSV function of the microchannels allowed either the central chamber, or the surrounding gyroid to be filled selectively with a solution without leakage to the other part, **Figure 5d-e**. The small footprint of the OoC system allowed us to 3D print 42 OoC chips in a single 35-min print run, demonstrating a high manufacturing throughput (**Figure S15, Supporting Information)**. 3D confocal images of cultured IMR-90 cells within the 3D-printed gyroid structure show a 3D organization of cells surrounding the central chamber (**Figure S16 and Video S2, Supporting Information**).

**Figure 5.**
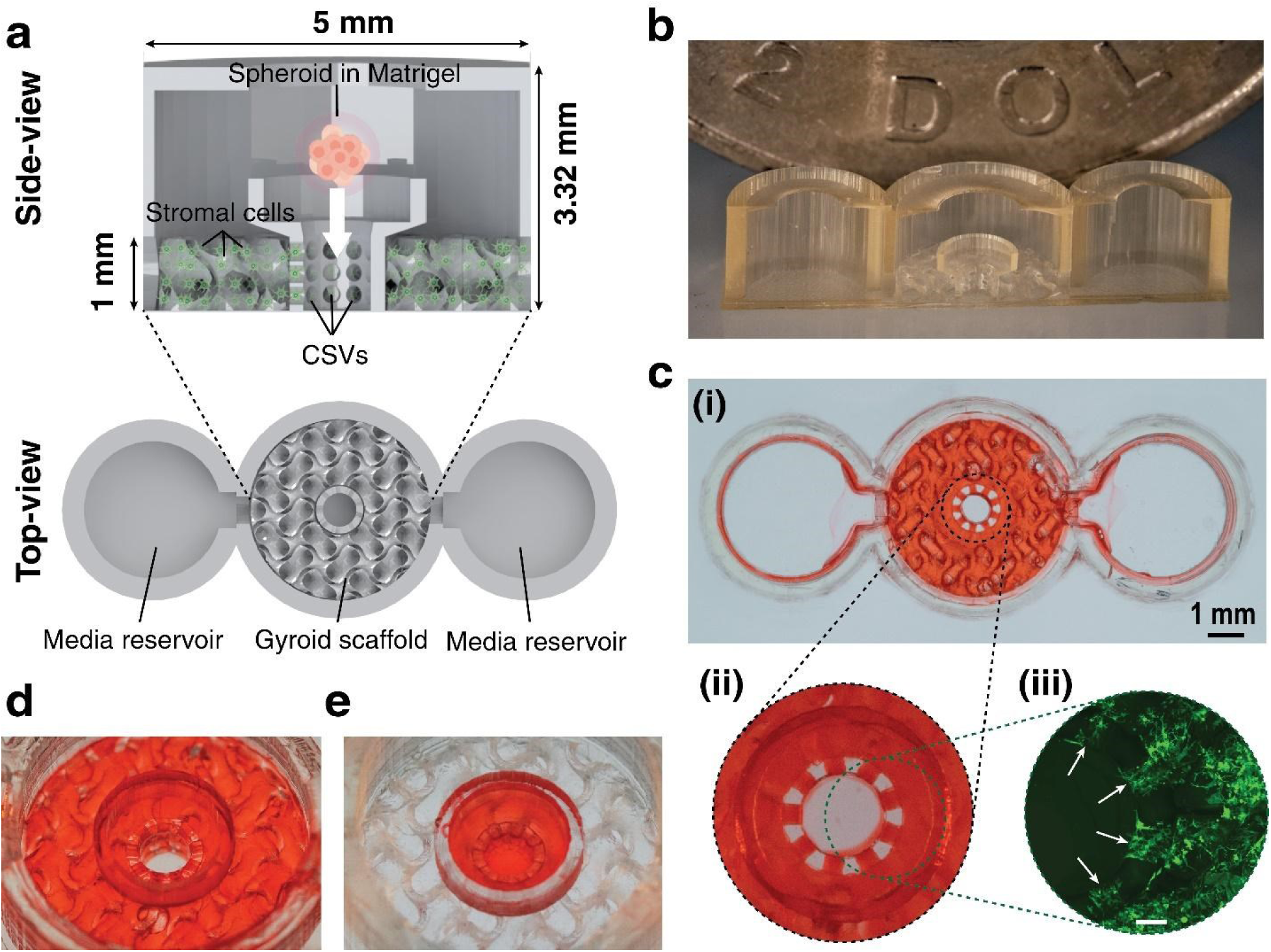
OoC device featuring CSVs and a gyroid scaffold. **(a-b)** Schematic representation of the OoC system layout that shows two reservoirs connected to the gyroid scaffold including the central openings, separation walls with a funnel structure, and microconduits on 3 levels connecting to the gyroid with CSVs on either side. **(b)** Close-up photograph of a 3D OoC system that was cut to show the internal structures with a Canadian $2 coin in the back. **(c) (i)** Top view of the OoC system filled with food dyed Matrigel except for the central chamber that is isolated by CSV. **(ii)** Zoomed-in view, highlighting the functionality of stop valves. **(iii)** showing the presence of stromal cells halted at the CSV (white arrows) (scale bar: 200 µm). **(d-e)** Illustration of the bidirectional functionality of CSVs during the seeding process.

To achieve three-dimensional seeding of stromal or endothelial cells throughout the entire gyroid structure in our OoC, we initially filled the two side media chambers with a cell-embedded Matrigel solution. The OoC device was kept on ice to prevent the Matrigel solution from gelating. The gyroid structure in the central chamber acted as a capillary pump, so the central chamber was spontaneously filled via capillary flow^36^, thus emptying the media reservoirs, and the surface tension of the liquid solution served to stop the self-filling at the multilayer CSVs. The device was removed from ice to allow the cell-embedded Matrigel solution to gelate within the central gyroid chamber. Subsequently, an MDA-MB-231 breast cancer spheroid was embedded in Matrigel and seeded in the central chamber. This assembly was then gelated in the cell incubator and filled with cell media, followed by bi-daily media changes and time-course imaging. The detailed OoC seeding protocol is described in the Method section. The cells on the gyroid were seeded throughout a height of 1 mm and could interface with the spheroid from all horizontal and vertical directions.

We initially assessed the long-term co-culturing performance of our OoC using a model cancer-stroma system comprising an MDA-MB-231 spheroid and the fibroblast cell line IMR-90. The cells were fluorescently labeled with tdTomato and GFP, enabling live-cell imaging throughout the time-course study. Over the 14-day period, both cell lines retained their physiological morphology within the device, as shown in **Figure 6a**. Notably, by day 8, we observed bidirectional migration of both cell types. Several IMR-90 single cells were seen migrating towards and into the spheroid chamber at different heights, and similarly, some interaction from the spheroid chamber was noticed (**Figure 6b**). Although it was initially unclear whether the MDA-MB-231 cancer spheroid migrated out of its chamber by day 8, we noted that by day 14, the bidirectional nature of cell migration was evident, with both cell types appearing to move towards each other.

**Figure 6.**
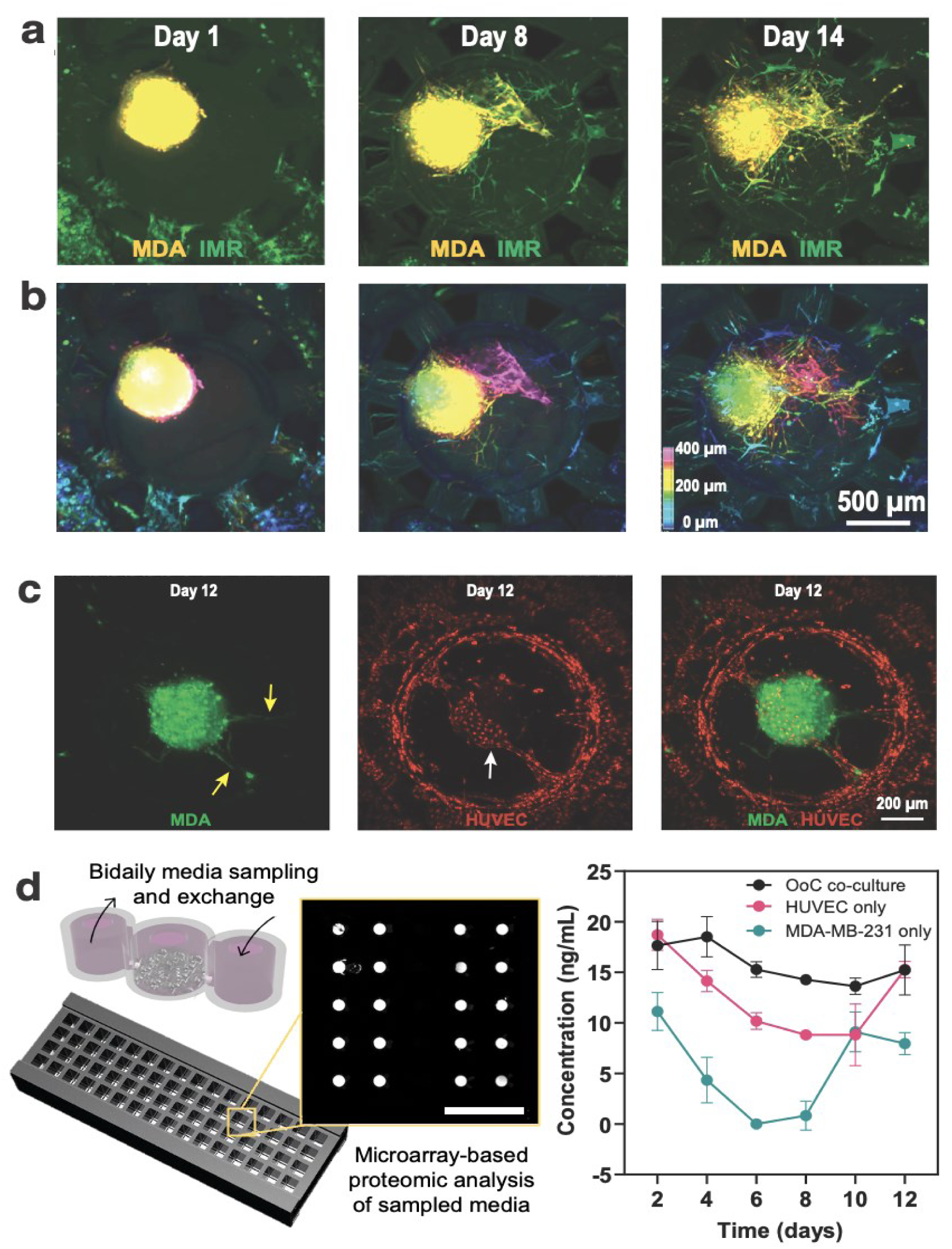
Co-culture of cancer and stromal cells in the OoC device. **(a)** 3D confocal microscopy images of tdTomato+ MDA-MB-231 spheroid co-cultured with GFP^+^ IMR-90 cells at different time points in the OoC platform. **(b)** Depth color map for the height range of 400 µm showing bi-directional migration of cells. **(c)** Confocal microscopy images of GFP+ MDA-MB-231 breast cancer cells spheroid co-cultured with mCherry+ HUVEC cells, demonstrating vessel formation and integration of the cancer spheroid into the vasculature (white and yellow arrows, respectively). **(d)** Schematic of microarray-based proteomic analysis and CXCL12 secretion under different culture conditions, showing increased concentration for OoC co-culture that induces cell migration. 3 biological repeats and 20 technical repeats were done for each condition and time course.

To further evaluate the utility of our OoC using a more sensitive cell type, we co-cultured green fluorescent protein positive (GFP+) MDA-MB-231 spheroids with mCherry+ HUVEC cells in the gyroid chamber. The co-culture was maintained for up to 12 days with media sampling and exchange every two days, followed by fluorescence confocal imaging at the endpoint (i.e., day 12). Additionally, mono-culture controls where either the cancer spheroid or the HUVEC cells were cultured alone in the OoC were performed, and media was also collected every two days. Similar to our previous co-culture setup, both HUVEC and MDA-MB-231 maintained their physiological morphology throughout the time course. We also observed bi-directional migration of both cell types, with HUVEC forming vessel-like structures surrounding the shell of the cancer spheroid (white arrow), and integration of the cancer spheroid into the vasculature (yellow arrow) **(Figure 6c)**. Furthermore, upon proteomic analysis of the time-course cell supernatant collected from the OoC, we found that the level of CXCL12 was consistently elevated throughout the duration of the study. Conversely, in both monocultures, the level of CXCL12 exhibited a decreasing trend until there was a sharp increase near the endpoint—day 12 for HUVEC and day 10 for the cancer cells **(Figure 6d)**. As CXCL12 has been associated with the promotion of directional cell migration. This pattern hints that chemotactic signaling was successfully maintained in the co-culture throughout the experimental timeframe, as indicated by the steady elevation of CXCL12. Meanwhile, cells in the monocultures initially down-regulated CXCL12, as evidenced by the decreasing levels of this signal. However, as the culture matured on the chip (i.e., closer to the endpoint), there was an upregulation of CXCL12 expression. Additionally, we monitored VEGF-A levels that were depleting in co-culture compared to the mono-culture of HUVECs, which would suggest a consumption in the OoC device^51^ (**Figure S17a, Supporting Information**). Meanwhile, we observed an increase in MMP9, which is associated with matrix breakdown and a prerequisite to cell migration^52,53^ (**Figure S17b, Supporting Information**). Based on these findings, we conclude that scaffolds made of P-PEGDA ink are well-suited for long-term studies involving organ-on-a-chip and 3D cell interactions.

## Discussion and Conclusion

In this study, we have successfully developed a high-resolution nanoporous PEGDA ink for fabricating OoC devices using VP 3D printing. 3D printing is a rapidly growing technology contributing to the advancement of microfluidics by facilitating the production of intricate and complex microstructures that are challenging to produce using conventional methods. We note a recent study on PIPS using methanol as a porogen and that was used to make thin membranes for DNA extraction^54^. This other study did not evaluate the potential for cell culture, structures were at a lower resolution and geometrically simpler compared to the ones shown here, and used a high concentration of BAPO photoinitiator (3%) which is likely cytotoxic. Notwithstanding these points, methanol might also not be a suitable porogen for making porous PEGDA suitable for cell culture and OoC applications. It remains challenging to 3D print such large-scale devices with high-resolution features owing to the trade-off between printing resolution, printing throughput, and overall built area. P-PEGDA overcomes this trade-off with an intrinsically nanoporous ink with a tunable pore size from 5-30 nm adjustable simply by varying the concentration of the porogen. With a vertical resolution of 27 µm, we were able to fabricate complex structures with precise control over the z-axis.

Our investigation into the nanoporous PEGDA ink reveals an interplay between porogen concentration, pore formation, gas permeability, and molecular diffusion. Despite a high porogen concentration of 30% and a corresponding mass loss of approximately 26%, the observed diffusion of molecules through the PEGDA matrix was slow, and concentration low, reaching only about 1.2% of the concentration in solution. This finding suggests that pore interconnectivity within the studied porogen range was limited, although still sufficient for oxygen gas permeability. Notably, P-PEGDA with a 10% porogen concentration exhibits oxygen permeability similar to that of PDMS, suggesting that P-PEGDA ink can effectively replace PDMS in applications requiring similar oxygen permeability. The oxygen permeability can be tuned based on the porogen concentrations for specific biomedical applications, particularly in organ-on-a-chip systems.

Our systematic cytotoxicity studies, conforming to ISO standards across four diverse cell lines, revealed high biocompatibility. Achieving over 80% cell viability emphasizes the potential of P-PEGDA for future *in-vitro* studies. The nanoporous substrates showed an increase in coverage by endothelial cells and a 77-fold increase for MDA-MB-231 cells, indicating strong cell attachment and improved cell interaction for *in vitro* studies.

In our study, we introduced P-PEGDA with 10-30% porogen as an easy-to-use photo-ink for 3D-printing meeting ISO standards for cell viability and promoting cell attachment. We proposed an ink that is suitable for biological applications and that can be 3D printed to meet the needs of any custom OoC design, and future work should aim to ideate new designs that recapitulate biologically relevant architectures and cell-level organization, vascularization, and alignment. Notably, our OoC system facilitates long-term cell interactions and bidirectional migration. Compared to conventional 2D microfluidic OoC designs which support cell interaction and migration only from beneath, this 3D OoC model supports interaction from all directions, and thus more closely resembling *in vivo* conditions. However, future work is needed to thoroughly examine and quantify the utility of 3D OoC in practical use cases, and in particular devise means of quantitatively assessing bidirectional migration through the OoC device. For example, the human bone is a naturally occurring triply periodic minimal surface (TPMS) structure, where its unique topological structure has been shown to regulate the migration and differentiation of bone-residing cells^55–57^. As such, many existing bone-on-a-chip models entail the enclosure of a decellularized bone scaffold or synthetic scaffolds inside a microfluidic device fabricated via PDMS molding or CNC micromachining^58–61^. Other OoC models have been limited to relatively planar devices, which reduces the surface area for crosstalk between adjacent reservoirs^43^. Furthermore, 3D printing has enabled scaffold fabrication directly within the OoC, which allows for a high surface area, and the potential for higher seeding density in a small footprint device, and one that promotes cell attachment, alignment, and gas permeability, which constitute the salient features of our P-PEGDA material^62^. In comparison, we have demonstrated P-PEGDA’s capability in manufacturing highly complex TPMS structures (i.e., gyroid), showcasing its potential in one-step fabrication of bone-on-a-chip models. Our current ink demonstrates the potential for 3D printed OoC designs; next steps towards validation with *in vivo* models should first aim to perform *in vitro* validation for a targeted OoC model, such as the aforementioned bone-on-a-chip designs. Beyond modeling organs with naturally occurring scaffold structures, we also speculate that 3D OoC models like ours may have a general advantage in promoting cell-cell interaction simply owing to the larger crosstalk interface between the cell chambers, given that 3D migration was observed in all three OoC cell types (i.e., IMR-90, MDA-MB-231, and HUVEC). Extending the design to multiple cell types can be foreseeable thanks to the customizability of 3D printing; while prior OoCs were limited to planar or multi-layered replica molded devices, here they can be designed to meet desired specifications and modified to contain multiple reservoirs for different cell types with CSVs and 3D-printed embedded channels to connect various reservoirs^36^. Media exchanges and perfusate can be studied downstream for proteomic analysis of compartmentalization and crosstalk. Challenges remain in sharing different culture conditions and media compositions, and for the integration of sensitive cell types (e.g., iPSCs), but advanced designs, multi-material capabilities, and customizability of 3D printing opens a myriad of possibilities.

Future work should also aim to extend the P-PEGDA formulation by considering smaller porogen sizes and higher monomer molecular weights. The current formulation using low-MW PEGDA-250 monomer and low-MW PEG-200 porogen limits the amount of porogen that can be added before failed layer adhesion. Modifications to the P-PEGDA formulation to broaden its applicability can include the use of a smaller porogen size such as methanol^54^, cyclohexanol^20^, or decanol^20^, so the porogen concentration can be increased while keeping a relatively low viscosity, and benefiting from increased porosity and pore size. Conversely, the monomer size can be increased by extending the P-PEGDA formulation towards higher molecular weight PEGDA-700, 1000, 3400 and 6000 for a reduced crosslinking density, reduced brittleness, and inherent permeability. This could expand the biomechanical and permeability properties of P-PEGDA over a tunable range, and potentially mimic other organ systems for the realization of 3D-printed multi-organ-on-a-chip models. Further studies will be needed to better understand the relationship between cell attachment, cell migration and scaffold characteristics such as porosity, pore size and connectivity, as well as benchmarking its performance in biologically relevant scenarios such as vascularization, tumor invasion, and tissue regeneration. The nanoporous P-PEGDA ink and the ease of 3D printing complex, high-resolution, gas permeable, and biocompatible scaffolds offer significant opportunities for OoC models, and might find use in drug discovery, drug response and regenerative medicine.

## Materials and Methods

### 3D printing

All objects were designed in SolidWorks® computer-aided drafting software, exported as “STL” files, and 3D-printed with MiiCraft Prime 110 (Creative CADworks, Concord, Canada) DLP 3D printer with a projected pixel size of 40 µm, featuring over 4 million pixels and a build area of 110 × 62 × 120 mm^3^, thus making it a suitable option for high resolution microfluidic 3D printing over a large build area. The 3D printer was equipped with a 385 nm illumination wavelength, which expanded material selection, and allowed for custom material 3D printing. All objects reported in this work are printed with a layer thickness of 20 µm, a single base layer exposure of 5 s, and normal layer exposure of 950 ms printed with a gap adjustment between the build plate and the bottom of the vat of 0.1 mm. Immediately after printing, to remove unpolymerized ink and clean channels, closed channels were vacuumed, washed with isopropanol (Fisher Scientific, Saint-Laurent, Quebec, Canada) several times using a syringe, and dried under a stream of pressurized nitrogen gas. All the 3D printing was done between 19-23°C at a relative humidity of 36-38%, which was monitored using a Traceable™ Jumbo Thermo-Humidity Meter (cat. #11-661-19, Fisher Scientific, Saint-Laurent, Quebec, Canada).

### Ink Formulation and Preparation

Inks in this study consist of a low molecular weight poly(ethylene glycol) diacrylate monomer (PEGDA-250, cat. #475629, Sigma-Aldrich, Oakville, Ontario, Canada), low molecular weight poly(ethylene glycol) porogen (PEG-200, cat. #P3015, Sigma-Aldrich, Oakville, Ontario, Canada), TPO photoinitiator (0.5% wt/wt) (cat. #415952, Sigma-Aldrich, Oakville, Ontario, Canada), and ITX photoabsorber (0.8% wt/wt) (cat. #I067825G, TCI America, Portland, Oregon, United States). The ink was prepared by massing the above-mentioned components using a digital precision analytical balance (XS204, Mettler-Toledo), then mixing under magnetic stir at room temperature for at least 30 min and up to overnight. Following their preparation, all inks are stored in amber glass bottles at room temperature for short-term storage and at 4°C for long-term storage until use.

### Videos and image stacking

Videos and images were taken using a Panasonic Lumix DMC-GH3K and Sony α7R III. For focus stacking, Imaging Edge Desktop (Sony) was used to take the sequence of images on different focal planes. Then, CombineZP was used to process the images. Imaging at the microscale was done on an inspection microscope (Nikon Eclipse LV100ND) at 5x magnification. Some images were taken with a stereo microscope (Leica SMZ-8) fitted with a digital camera (Lumix GH3 DSLR, Panasonic).

### Penetration depth measurements

In order to measure the effect of porogen on the penetration depth of light, a 5 µL drop of the formulation was placed on a glass slide that was silanized by liquid phase deposition in a solution of 2% 3-(trimethoxysilyl)propyl methacrylate (cat. #M6514, Sigma-Aldrich, Oakville, Ontario, Canada) prepared in toluene (cat. #T3244, Fisher Scientific, Saint-Laurent, Quebec, Canada) for at least 2 h or overnight. The slides were cleaned with fresh toluene and dried under a stream of compressed nitrogen gas, then placed directly on the 3D printer light source. The power intensity of the 3D printer light engine was measured through the glass slide using a UV light intensity meter with a 385-nm-illumination probe (Model 222, G&R Labs, Santa Carla, California, United States) to be 5 mW/cm^2^, and the formulations were then exposed to 385 nm illumination at varied exposure times. The subsequent energy dosage was given as the product of the illumination irradiance and the exposure time. After photocuring, the slide was cleaned with 70% ethanol to remove excess uncured ink. The cured ink height was then measured using a stylus profilometer (DektakXT, Bruker Co.). To complement the penetration depth measurement, the absorbance of all the formulations was measured in triplicate using a NanoDrop spectrophotometer (ND-1000, NanoDrop Technologies, Wilmington, Delaware, United States). The absorbance spectra of all the ink formulations was recorded using 2 µL of the ink at a 0.1 mm path length for high absorbance samples.

### Viscosity measurement

The viscosity of the P-PEGDA inks was measured using a vibrating rod viscometer (SV-10, A&D Company). The viscometer receptacle was loaded with 50 mL of the formulations and the sensing rods were lowered into the receptacle until they were fully immersed in the ink. Then, the sensing rods were set to vibrate at a frequency of 30 Hz to measure the flow resistance. All measurements were done from 19-20°C.

### SEM

The Hitachi SU-8230 SEM was used to characterize the pore structure of the 3D-printed objects at an operating voltage of 3 kV. Before the SEM measurements, the samples were washed, dried, and then coated with a 4 nm thick platinum layer. Pore sizes were measured from the cross-sectional SEM images by measuring 40 random pores in ImageJ.

### Atomic force microscopy (AFM) measurements

The surface roughness of 3D printed P-PEGDA was measured by AFM using a Multimode 8 AFM (Bruker Co., Billerica, Massachusetts, United States). For each ink formulation, an 8 × 8 × 3 mm^3^ (length × width × height) sample was 3D printed, washed in ethanol for 48 hours with daily refresh, dried with compressed nitrogen, and then mounted onto the AFM stage. A 10 × 10 µm^2^ region was scanned using a triangular tip ScanAsyst-Air cantilever probe with a nominal tip radius of 2 nm, spring constant of 0.4 N/m, and tip length of 115 µm. The scans were analyzed in Gwyddion (version 2.59, public domain software, Czech Metrology Institute) to generate 2D and 3D topology profiles and determine the mean surface roughness.

### Mass loss

Disk-shaped samples with a diameter of 8 mm and a thickness of 3 mm were 3D-printed and washed briefly with 70% ethanol to remove uncured ink. The samples were dried with compressed nitrogen and then massed on a digital precision analytical balance (XS204, Mettler-Toledo). Then, the samples were immersed in 1X PBS and shaken on an orbital shaker for 48 h for porogen removal. The samples were dried under compressed nitrogen and massed again to determine the mass loss due to porogen removal. The difference in mass normalized by the initial mass was used to determine the mass loss ratio.

### Contact angle measurements

A 2 µL drop of deionized water was placed on the top surface of 3D-printed samples and imaged using a Panasonic Lumix DMC-GH3K. The side view images were imported to ImageJ to measure the static contact angle using the contact angle plugin. Measurements were taken between 19-23°C at a relative humidity of 36-38% for all 3D-printed samples, and monitored using a Traceable™ Jumbo Thermo-Humidity Meter (cat. #11-661-19, Fisher Scientific, Saint-Laurent, Quebec, Canada).

### Gas permeability assay

To test for the degree of oxygen permeability in our 3D printed P-PEGDA formulations, we employed an oxygen sensing test strip by dip-coating cellulose paper with 500 mg/L resazurin solution, where the test strip will change color when oxygen level changes. We set up gas impermeable rubber tubes containing the test strip and capped it with barrier discs of either no barrier, an impermeable aluminum barrier, 1:10 PDMS (Sylgard 184 Clear Kit, cat. #MSPP-DC2065622, Krayden, Canada) discs, or 3D printed P-PEGDA discs with varying porosity. The discs were 8 mm in diameter and 3 mm thick, while the flexible tube had an inner diameter of 7 mm, an outer diameter of 9 mm, and a height of 30 mm. The tubes were put under vacuum, where the only outlet for gas to escape is through the barrier cap. Uncapped test strips were placed under the same vacuum condition and its color readout was defined as 100%, while test strips placed under normal oxygen condition (i.e., atmospheric) was used to define 0%. The experiment was until obvious color change is observed in the uncapped, vacuumed test strip condition (∼48 hours) and the color of each test strips was quantified using Fiji (ImageJ). P-PEGDA 3D printed discs were measured in 2 independent experiments from 2 separate print batches. PDMS and impermeable aluminum barrier controls were measured with 1 repeat only, so the error bar represents the standard deviation in the quantified color readout within the test strip region. In all other conditions, the error bar represents the standard deviation between the 2 independent repeats.

### Data analysis

The error bars displayed in the figures represent the standard deviation. GraphPad Prism 9 was used for data analysis.

### Cell culture

The normal human fibroblast cell line IMR-90 (ATCC CCL-186), triple-negative breast cancer cell line MDA-MB-231, and immortalized kidney cell line 293T were cultured in Dulbecco’s Modified Eagle Medium (Gibco, USA) containing 4.5 g/L D-glucose, 4.5 g/L L-glutamine, and 110 mg/mL sodium pyruvate. The media was further supplemented with 10% FBS and 1% penicillin-streptomycin (Gibco, USA). Fluorescently tagged MDA-MB-231 and IMR-90 were cultured under the same condition as the wildtype cell line. Human umbilical vein endothelial cells (HUVEC) expressing the fluorescent protein mCherry were kindly provided by Dr. Arnold

Hayer of McGill University^63^. HUVEC cells were cultured in EGM-2 media (Lonza, USA). All cell lines were incubated at 37°C with 5% CO_2_ supplementation. The cells were grown and passaged according to ATCC’s recommendations. HUVECs were used at passage 6 and all other cell lines (i.e., 293T, MDA-MB-231, IMR-90) were used between passage 15-20; in all cases, all biological repeats were done with cells from the same passage or the subsequent passage.

### Live/dead cell viability and cell attachment

Cell viability was calculated via live/dead signal quantification and performed using the LIVE/DEAD™ Viability/Cytotoxicity Kit (cat. #L3224, Invitrogen, USA). Briefly, ∼10k IMR-90 cells were seeded in 96-well plates and co-cultured with 8 × 3 mm^2^ (diameter × height) 3D-printed rings with either no post-treatment or with 12 h EtOH washing. PDMS rings of the same size were used a positive control. At 48 h, the cells were stained with Calcein-AM and Ethidium homodimer-1 for 30 min at 37°C to stain for live and dead cells, respectively. The entire microwell was imaged and signal of the Calcien-AM was used to create a thresholding mask in ImageJ to measure the cell surface coverage. Cell viability was calculated by setting the cell coverage reported by PDMS control as 100%. The cell attachment quantification was similarly calculated by normalizing the surface coverage of fluorescently tagged HUVEC and MDA-MB-231 cells on the various 3D-printed disks to the total area imaged.

### OoC seeding and maintenance

The media reservoirs were first loaded with a cell-embedded Matrigel solution consisting of ∼50,000 cells suspended in ∼20 µL of 25% Matrigel diluted in media and placed on ice to prevent the Matrigel solution from prematurely gelating. The solution self-filled the central gyroid chamber via capillary flow and stopped at the CSVs. Following the completion of capillary filling, the scaffold was incubated for 2 h in a cell incubator and flipped upside-down after 1h to promote uniform cell attachment. Subsequently, a three-day-old MDA-MB-231 breast cancer spheroid, previously cultured in a 96-well ultra-low attachment plate with an initial seeding density of 100,000 cells per well, was embedded in ∼10 µL 25% Matrigel DMEM solution and placed in the central chamber. The OoC was placed on ice while the spheroid was allowed to sediment to the bottom of the central chamber. This assembly was then gelated in the cell incubator for 30 minutes. Finally, the OoC was filled with approximately 200 µL of cell media, followed by bi-daily half-media changes (i.e., 100 µL simultaneous removal and addition of media) for proteomic analysis followed by time-course imaging through the OoC device. Moist tissues wetted with sterile water were placed within the well plates containing the OoC to prevent evaporation of the cell- and spheroid-embedded Matrigel solutions throughout the seeding process.

### Protein Microarray for CXCL12 Quantification

The CXCL12 antibody microarray was fabricated onto 2D-Aldehyde glass slides (PolyAn, Germany) using a sciFLEXARRAYER SX microarray spotter (Scienion) equipped with a single piezo dispense capillary nozzle with coating 1 (Scienion). Anti-CXCL12 antibodies (Cat# MAB350, R&D Systems) were diluted in spotting buffer containing 15% 2,3-butanediol and 1 M betaine in PBS to achieve a final concentration of 100 μg/mL. A relative humidity of 65% was maintained throughout the fabrication process and the printed microarrays were incubated overnight in the dark in a 70% humidity bell jar. A 5-dot by 5-dot microarray was fabricated consisting of 20 dots of anti-CXCL12 antibody and 5 dots of negative control antibody. 64-well ProPlate® multi-well chambers (Grace Bio-Labs) were used to enclose the fabricated microarray. Supernatant of 6 time-points collected from both the MDA-HUVEC co-culture and the respective mono-cultures were diluted 1:4 in PBS with 0.1% Tween-20 and 3% BSA. CXCL12 antigens with a concentration range of 78.125 pg/mL to 20 ng/mL were diluted in the same buffer as the experimental samples (i.e. 1-part EGM-2 media, 4-part dilution buffer) to allow quantification of protein concentration. The fabricated microarray slides were blocked in PBS 0.1% Tween-20 3% BSA for 2 h on the shaker at room temperature. Experimental sample and standard curve spike-ins were incubated on the array overnight at 4°C. Biotinylated anti-CXCL12 antibody (Cat# BAF310, R&D Systems) at the working concentration of 2 µg/mL was added to the microarray and incubated for 2 h at room temperature, followed by detection via Alexa647-conjugated streptavidin for 1h at room temperature. Finally, the microarray slide was imaged using an InnoScan 1100 AL microarray scanner (Innopsys) and the relative fluorescence unit of each spot were quantified using the ArrayPro 4.5 Software (Meyer Instruments).

### Cytotoxicity Testing

The cytocompatibility assays were performed in compliance with International Organization for Standardization (ISO) standards for the development of medical devices. Cells were seeded in a 24-well plate at a seeding density of 10,000 cells per well. Immediately after cell seeding, post-treated disks were placed directly on top of the cell layer. The 3D-printed discs were 8 mm in diameter and 3 mm thick. As controls, cell-only wells were also seeded with the same cell density. Quantitative cell viability measurements via PrestoBlue™ (Thermofisher, USA) were taken every 24 h for a time course of three days. Microscopy images were taken daily using a Ti2 inverted microscope and analyzed using NIS-Element (Nikon, Japan) in order to monitor cell morphology. Three biological replicates were done for each of the three above-mentioned conditions. For the PrestoBlue™ cell viability assay measurement, two technical replicates were performed for each biological repeat. In addition, the media color was monitored daily.

### Diffusion assay

The permeability of the P-PEGDA ink was evaluated by a confocal fluorescence microscope. Posts of 500 μm diameter and 5 mm height were 3D printed and immobilized in an observation chamber. The pinhole of the confocal microscope was set to 1.2 Airy Unit and a single plane at around 200 μm above the base of the post was imaged. Then, a 10 μM fluorescein solution in MilliQ was introduced within the chamber, and fluorescence images were acquired every minute, while keeping the shutter off between measurements to prevent photobleaching of the dye. By imaging a section away from the base and the top of the 3D-printed posts, the radial diffusion of fluorescein in the P-PEGDA is expected to account for increasing fluorescence intensity during the time of the experiment. Images were analyzed by a custom MATLAB script, where the center of the post was automatically detected via the MATLAB function *imfindcircle*. A circular mask was generated and separated into 9 distinct sectors. These sectors were individually applied onto the fluorescence images acquired in time to assess the homogeneity of the radial diffusion of fluorescein within the post. A second integration mask was generated to measure the fluorescence intensity outside the post and used to normalize the fluorescence intensity and account for photobleaching. In analogous to fluorescence recovery after photobleaching experiment (FRAP), and due to the radial diffusion of fluorescent molecules into a circular non-fluorescent material, a non-linear least-square function was fitted to the normalized intensity. The fit-function was 𝑓(𝑡) = 𝐴(1 −𝑒𝑥𝑝 𝑒𝑥𝑝 (−𝜆𝑡) ) + 𝑥_𝑜_ where A and λ are fit parameters and x_0_ the initial fluorescence intensity into the post.

## Acknowledgments

**(a)** V. K. acknowledges FRQNT for the doctoral scholarship (no. 268838). D. J. acknowledges support from NSERC strategic grant (no. 506689-17). D.J. acknowledges support from a Canada Research Chair in Bioengineering.

