## Supplementary Information for "Nanoporous PEGDA ink for High-Resolution Additive Manufacturing of Scaffolds for Organ-on-a-Chip"

Figure S1. Solubility of LAP in PEGDA-250

Figure S2. Comparison of transparency

Figure S3. Impact of porogen on ink absorption

Figure S4. Viscosity of P-PEGDA formulations

Figure S5. 3D-printed 3Dbenchy model at different porogen concentrations

Figure S6. Surface profiles of P-PEGDA formulsations

Figure S7. Assessing fluorescence intensity inside the P-PEGDA at various porogen concentrations

Figure S8. Gas permeability of P-PEGDA formulations

Figure S9. Effect of PA on cell viability

Figure S10. A comparison of the cytotoxicity between the P-PEGDA ink and previously published low molecular weight PEGDA-based inks

Figure S11. Fluorescence images of HUVEC cells cultured within enclosed channels at various time points

Figure S12. Cell viability of MDA-MB-231 with different porogen concentrations

Figure S13. Live/dead cell staining microscopy images

Figure S14. Quantitative cell alignment of HUVECs on P-PEGDA

Figure S15. 3D printing throughput for the OoC Device

Figure S16. Cultured IMR-90 cells within the 3D-printed gyroid structure

Figure S17. Proteomic profile of VEGF-A and MMP9 in the OoC device

**Supplementary Figures**

**Figure S1**

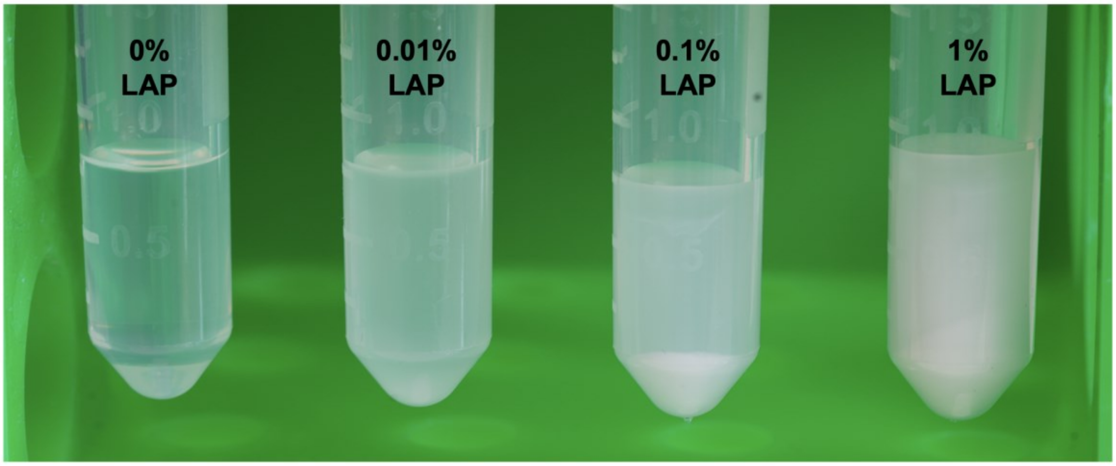


**Figure S1. Solubility of LAP in PEGDA-250.** LAP photoinitiator was added to PEGDA-250 and mixed for 30 mins to assess its solubility; cloudy solutions with visual sedimentation suggested poor solubility.

**Figure S2**

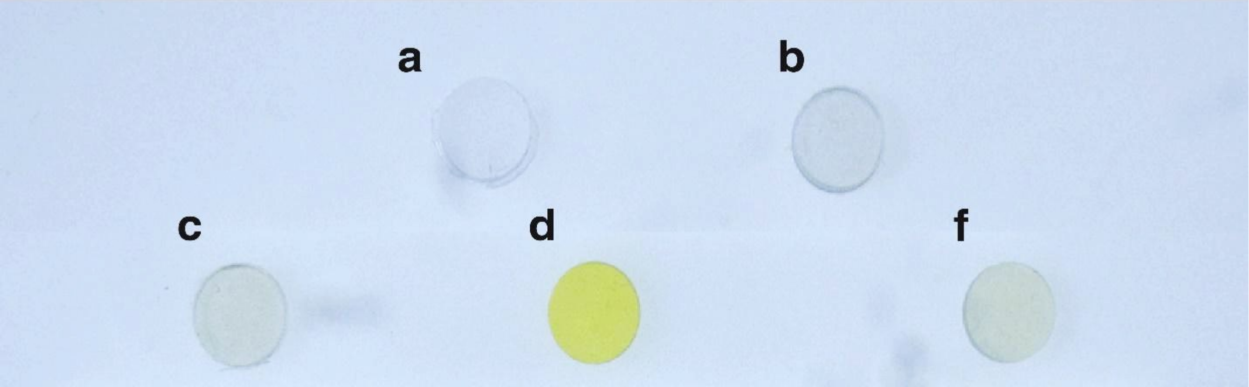


**Figure S2. Comparison of transparency in our formulation versus previously published formulations.** 3D-printed disks were prepared with different inks: **(a)** PEGDA+0.4% TPO, **(b)** PEGDA+0.4%TPO+0.8%ITX, **(c)** PEGDA+0.5%BAPO+0.8%ITX, **(d)** PEGDA+1%BAPO+2%NPS, and **(f)** PEGDA+1%BAPO+0.38%Avobenzone. Our formulation incorporates TPO, making it less toxic and less yellowish compared to previously published PEGDA formulations by Kuo et al.[1] **(c)**, Noriega et al.[2] **(d)**, and Warr et al.[3] **(f)**.

**Figure S3**

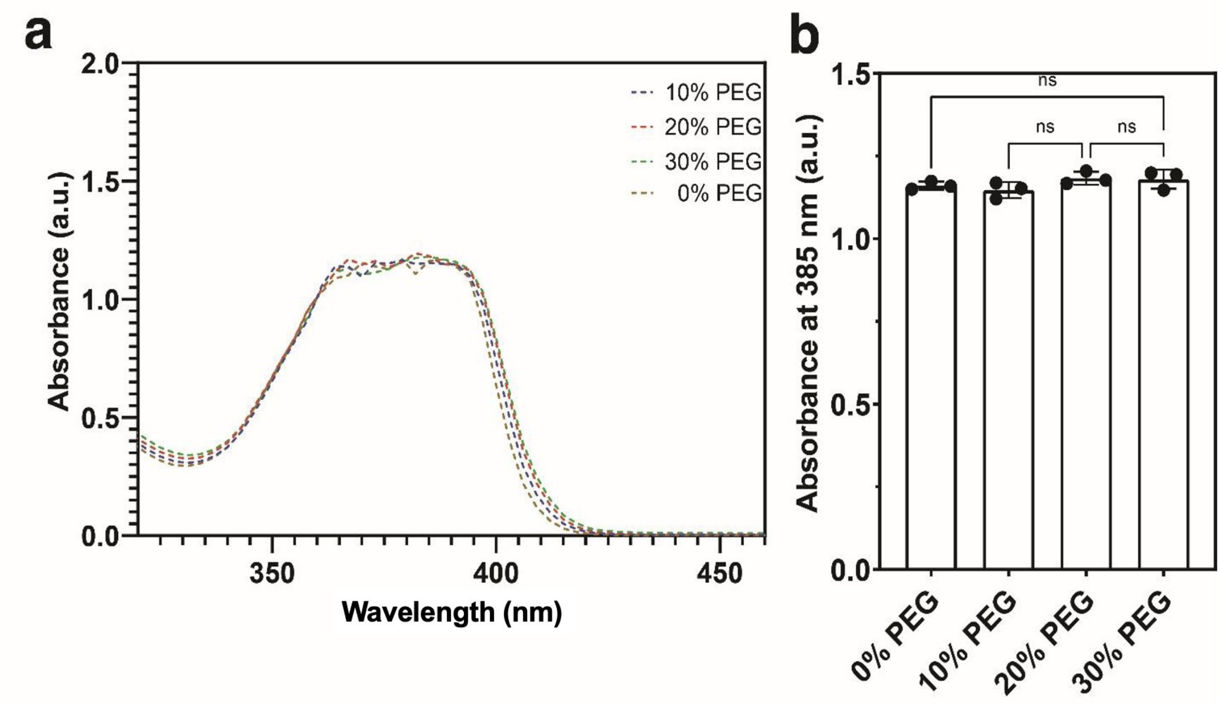


**Figure S3. Impact of porogen on ink absorption at 385 nm illumination wavelength of DLP projector. (a)** Absorption of P-PEGDA for varying PEG concentrations. **(b)** Absorption of P-PEGDA at 385 nm. Porogen concentration does not affect the light absorption of the ink. Mean values of N=3 measurements and error bars are the standard deviation. This is due to the low absorption of PEG porogen compared to PI and PA.

**Figure S4**

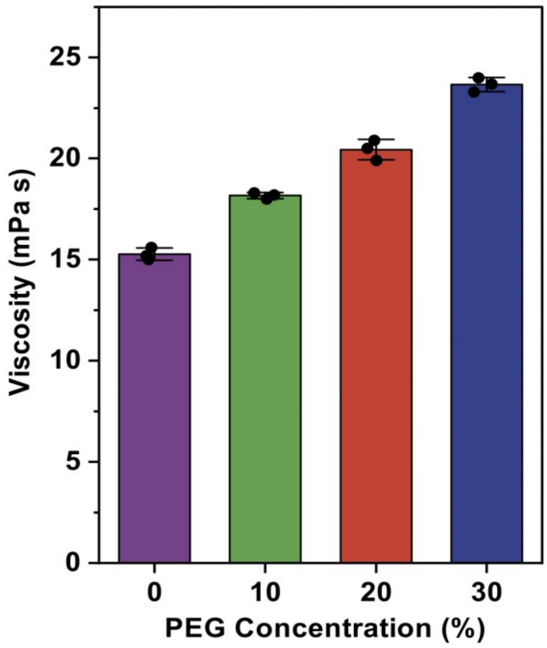


**Figure S4. Viscosity of P-PEGDA formulations.** Higher porogen concentrations slightly increases the ink viscosity but remain sufficiently low for high-resolution 3D printing.

**Figure S5**

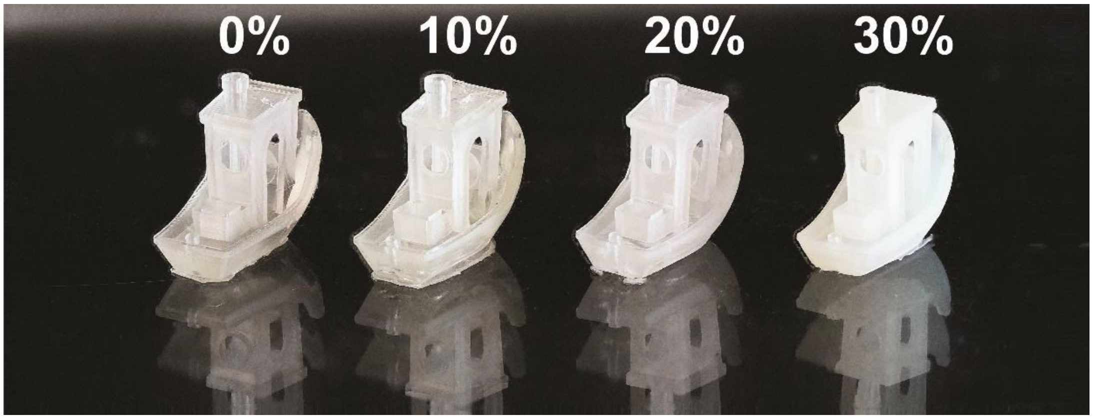


**Figure S5. 3D-printed 3Dbenchy model at different porogen concentrations.** Higher porogen concentrations result in decreased transparency of the 3D-printed parts.

**Figure S6**

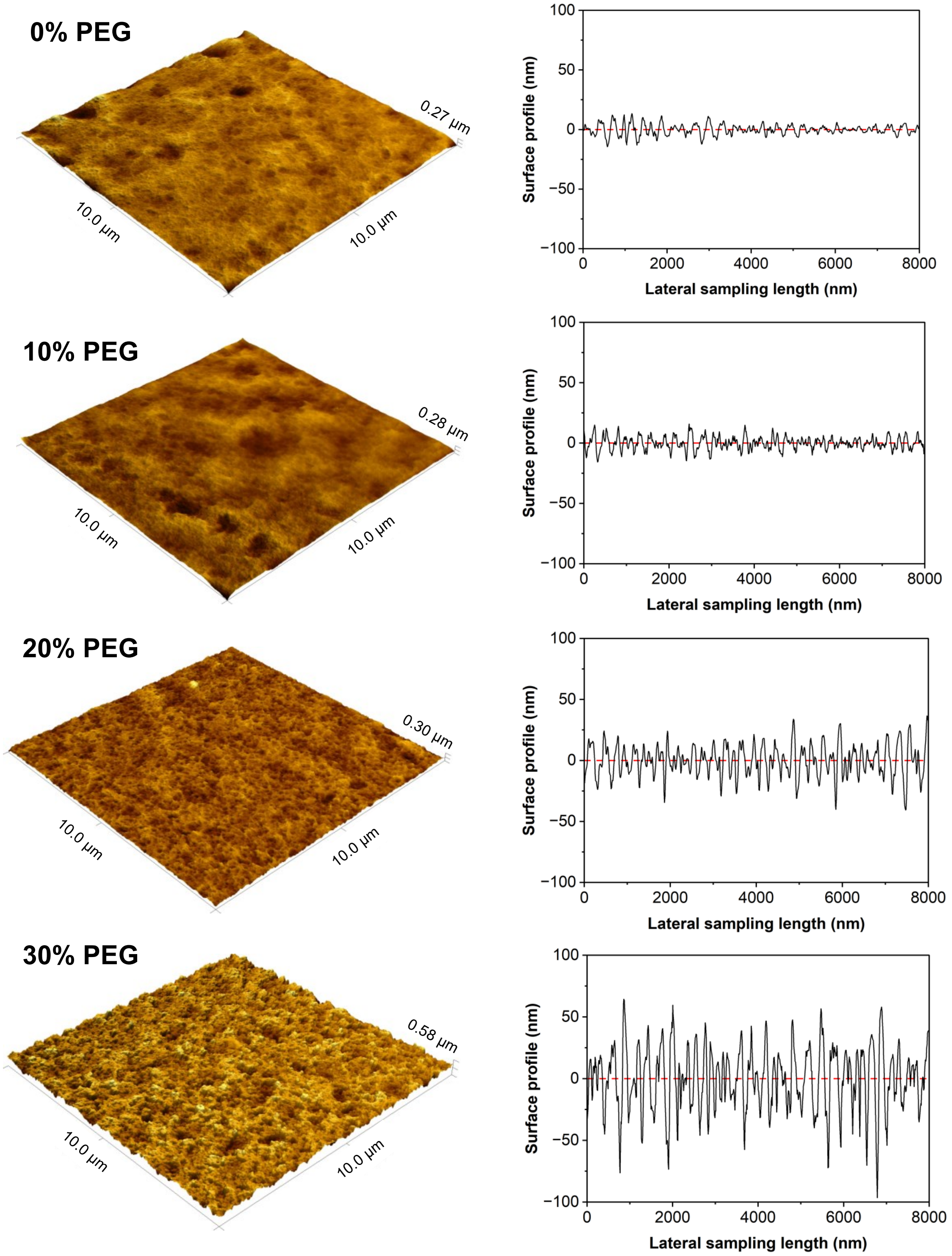


**Figure S6. Surface profiles of P-PEGDA formulations.** AFM surface profiles show that higher porogen concentrations lead to a greater surface roughness.

**Figure S7**

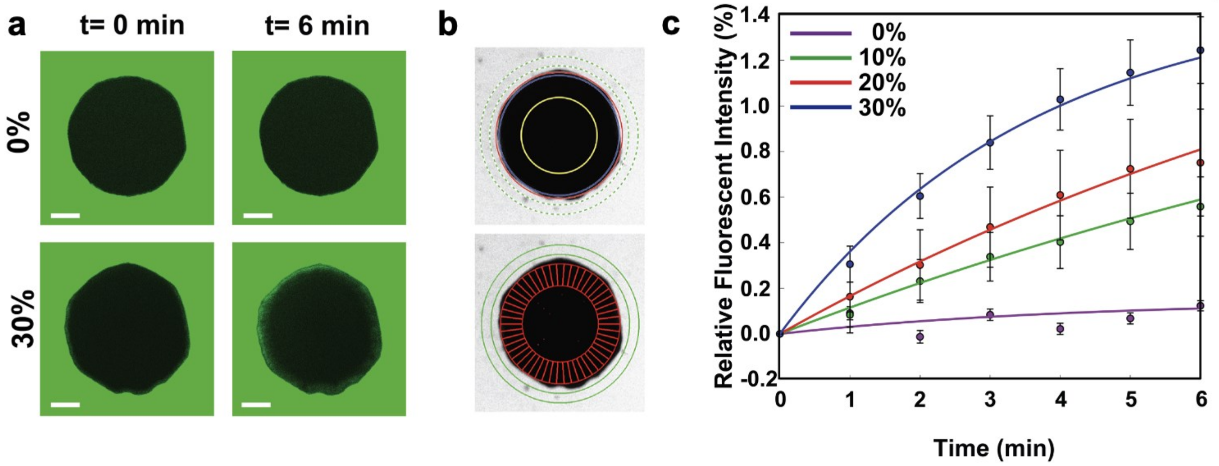


**Figure S7. Assessing fluorescence intensity inside the P-PEGDA at various porogen concentrations. (a)** Fluorescence confocal microscopy images of 3D-printed rods fabricated without porogen (top) and with 30% porogen (bottom). The images track the fluorescent dye as it diffuses into the rods. **(b)** Rod imaged at the mid-plane, where the rod is automatically detected by thresholding. The centroid of the rod is determined by its location, and its radius is determined as well. (red circle). To minimize background interference, we applied a 0.95 scaler to the radius, to generate an inner mask (blue circle). Another scaler (0.60) is applied to generate the inner boundary of the mask (yellow circle). The area in between the blue and yellow circle is then radially segregated into sectors to assess fluorescence across the periphery of the rods, and account for non-uniformity of the 3D-printed shape. In addition, we generated a disk-shaped mask (green circles), which is used to evaluate the mean fluorescence intensity of the bulk and used to normalize the signal measured inside the rods. **(c)** Time-course of the relative fluorescent intensity of fluorescein (340 g/mol) over time through the 3D printed rods with different porogen concentrations (See Figure S4, Supporting Information). Curves are non-linear least-square regressions (see method). P-PEGDA 3D-printed rods are immobilized in fluidic chamber and imaged by confocal microscopy. The outer phase is supplemented with 10 μM fluorescein in PBS pH 7.4.

**Figure S8**

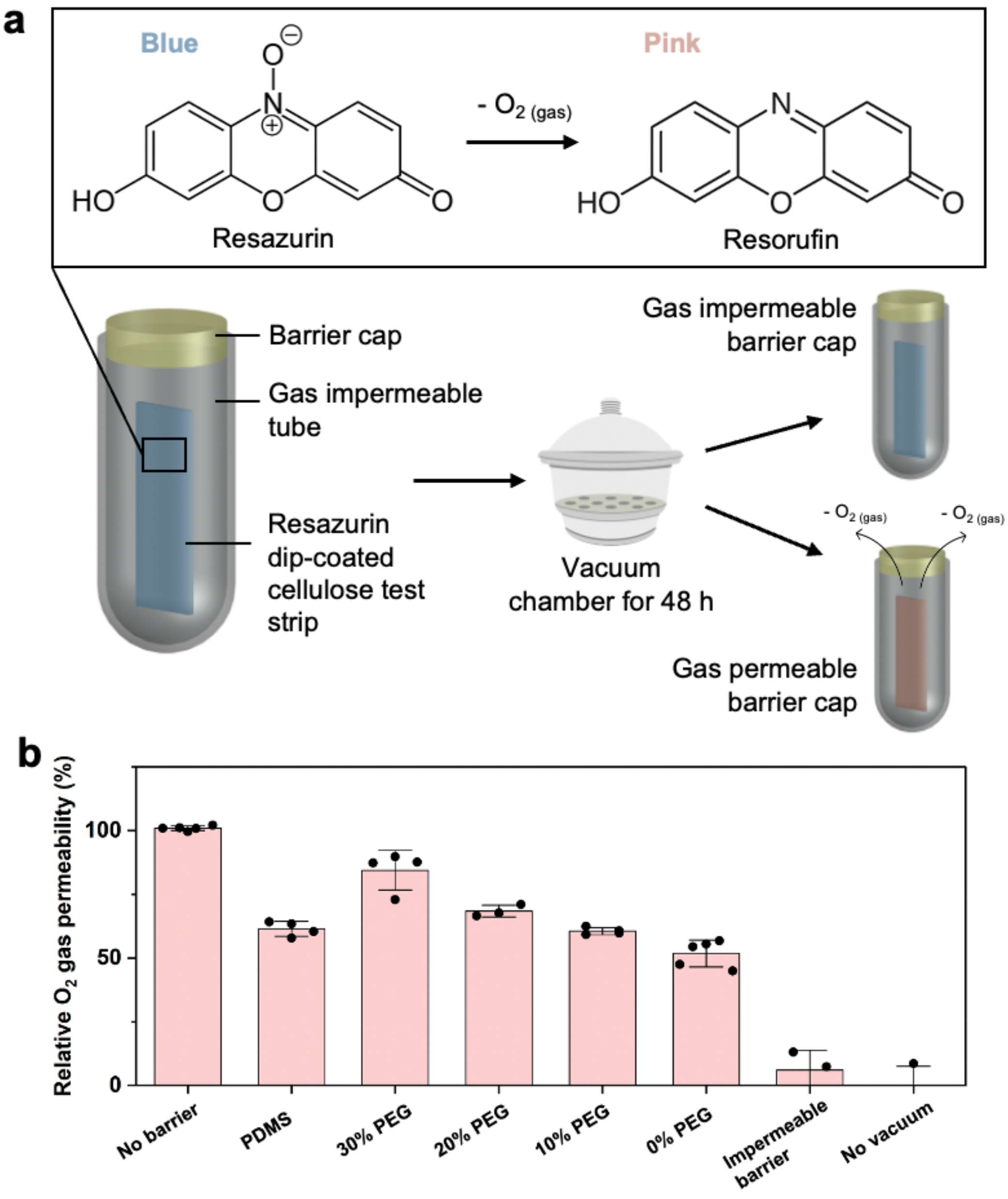


**Figure S8. Gas permeability of P-PEGDA formulations. (a)** Oxygen gas permeability measured by the reduction of resazurin on cellulose test strips enclosed in a gas impermeable tube capped with either no barrier, an impermeable aluminum barrier, PDMS (1:10 Sylgard 184) discs, or 3D printed P-PEGDA discs with varying porosity. After applying a vacuum, oxygen can exit the gas permeable barrier caps and result in the reduction of resazurin to resorufin (blue to pink color change), while the gas impermeable test strips retain the oxygen gas in the tube and undergo no color change. **(b)** Percentage of pink color change on resazurin dip-coated cellulose test strips.

**Figure S9**

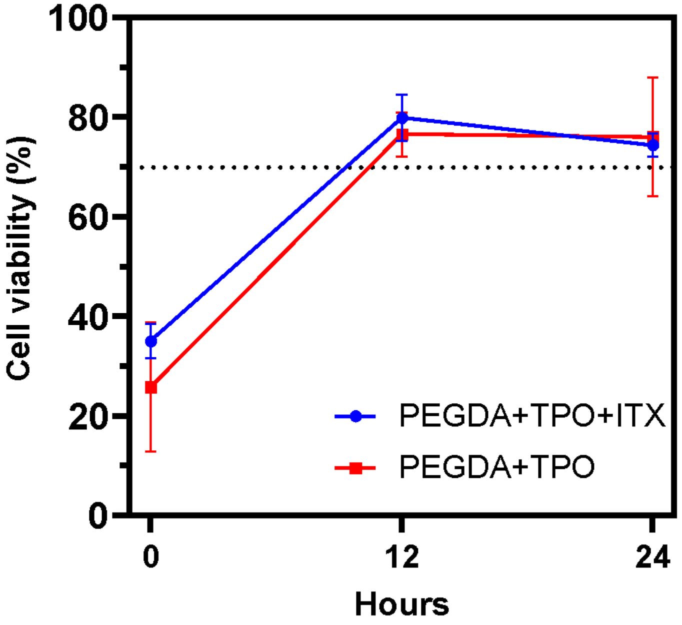


**Figure S9. Cell viability of 293T cells was analyzed using the ISO 10993-5 standard protocol for P-PEGDA ink with and without ITX PA, as a function of wash time.** The results show low toxicity of the PA and enhanced biocompatibility for washed samples.

**Figure S10**

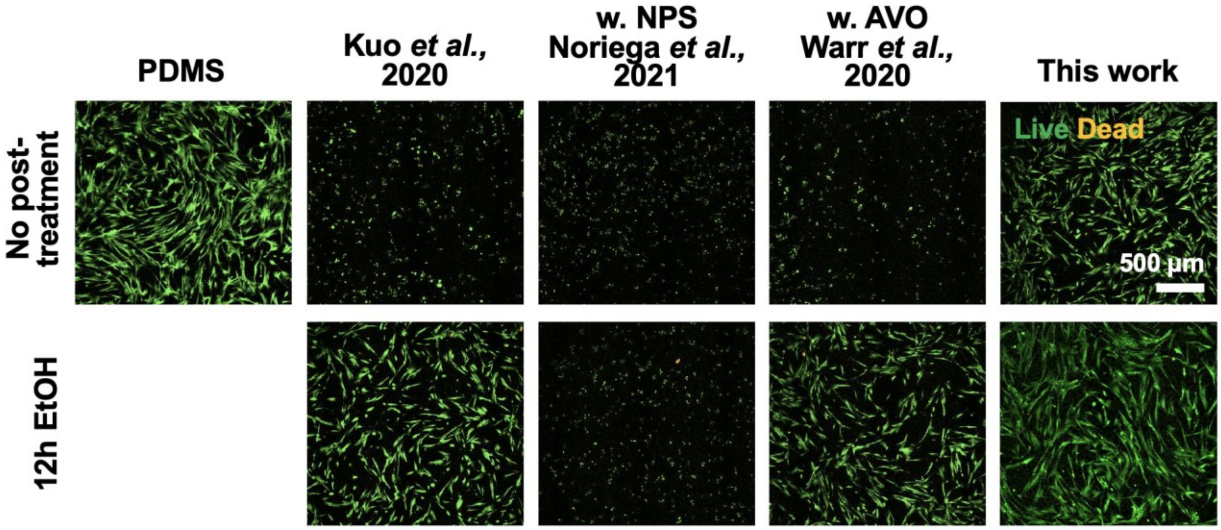


**Figure S10. A comparison of the cytotoxicity between the P-PEGDA ink and previously published low molecular weight PEGDA-based inks.** Fluorescence images of IMR-90 cells after 48 h of culture in a well-plate. The images compare samples that underwent a 12 h wash with 70% EtOH post-printing with those that received no wash. Our P-PEGDA formulation exhibits superior biocompatibility compared to the PEGDA formulations previously published by Kuo et al.[1] (containing 0.4 % BAPO PI and 0.8 % ITX PA), Noriega et al.[2] (containing 1% BAPO PI and 2% NPS PA), and Warr et al.[3] (containing 1% BAPO and 0.38 % Avobenzone), even without washing. Our P-PEGDA formulation, which uses TPO PA instead of the more cytotoxic BAPO, showed a higher cell viability.

**Figure S11**

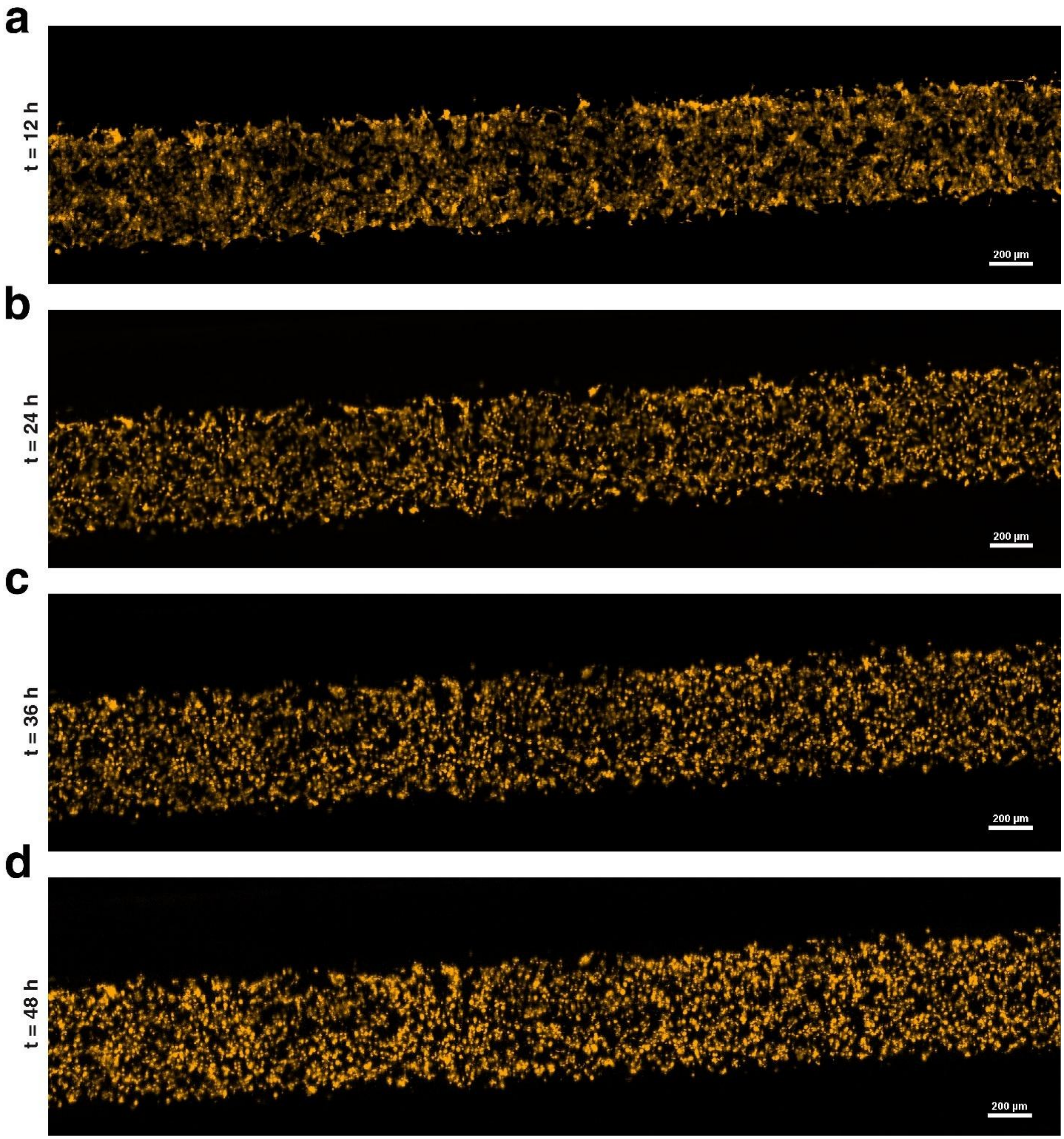


**Figure S11. Fluorescence images of HUVEC-mCherry cells cultured within enclosed channels at various time points: (a)** 12 h, **(b)** 24 h, **(c)** 36 h, and **(d)** 48 h. The device was enclosed with a luer-lock cap and kept inside a live cell chamber for continuous time-course imaging (every hour for 48 h). These images demonstrate the feasibility of culturing and imaging cells within an enclosed device for continuous monitoring.

**Figure S12**

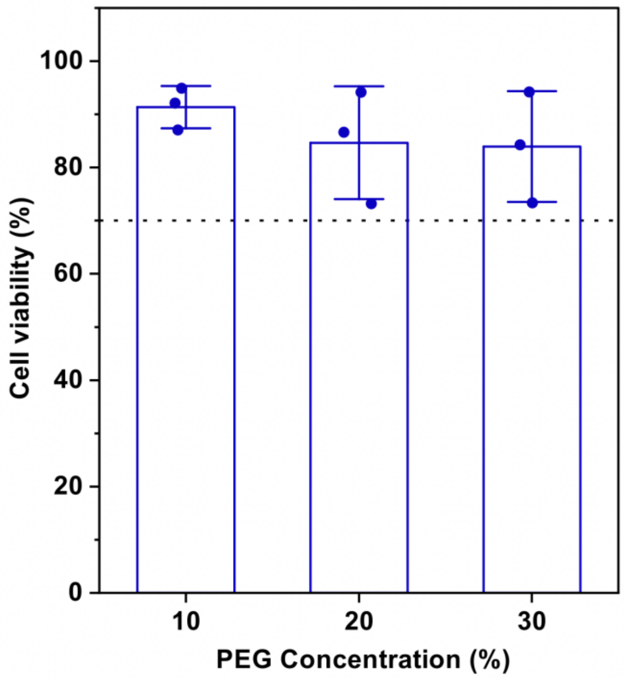


**Figure S12. Cell viability of MDA-MB-231 with different porogen concentrations.** Despite the addition of PEG porogen, the cell viability remains relatively unchanged.

**Figure S13**

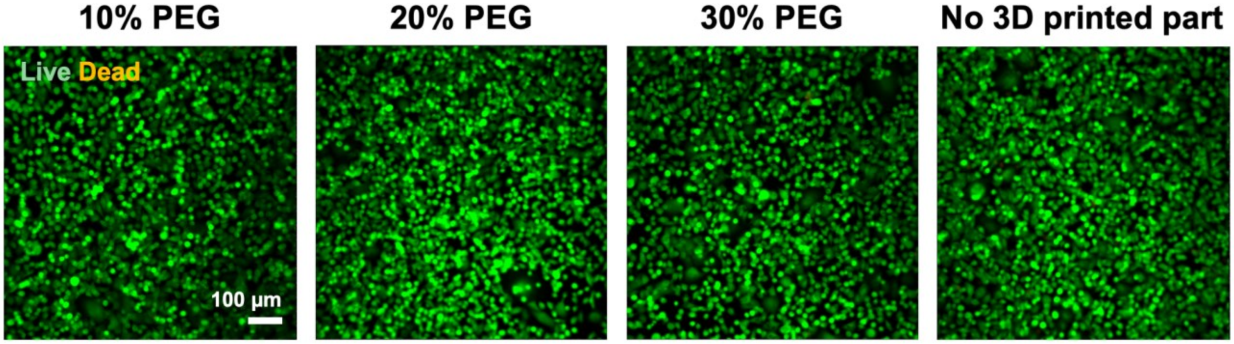


**Figure S13. Live/dead cell staining microscopy images.** Microscopy images showing live/dead cell staining following 72 hours co-culture of the 3D printed part with MDA-MB-231 cells.

**Figure S14**

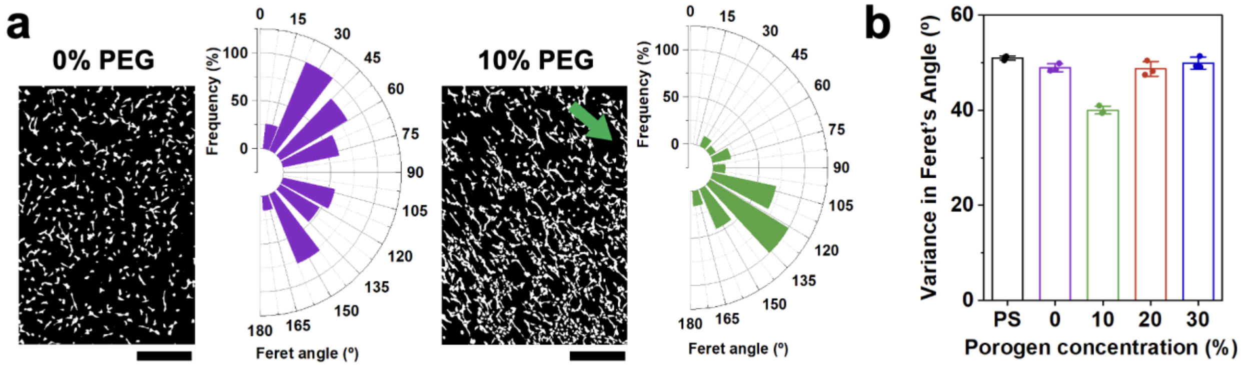


**Figure S14. Quantitative cell alignment of HUVECs on P-PEGDA. (a)** Measurement of Feret's angle of HUVECs on 3D printed P-PEGDA containing 0% and 10% PEG porogen. Scale bars = 500 µm. **(b)** Variance in cell alignment for P-PEGDA with different porogen concentrations, showing the least difference in Feret's angle at 10% porogen concentration.

**Figure S15**

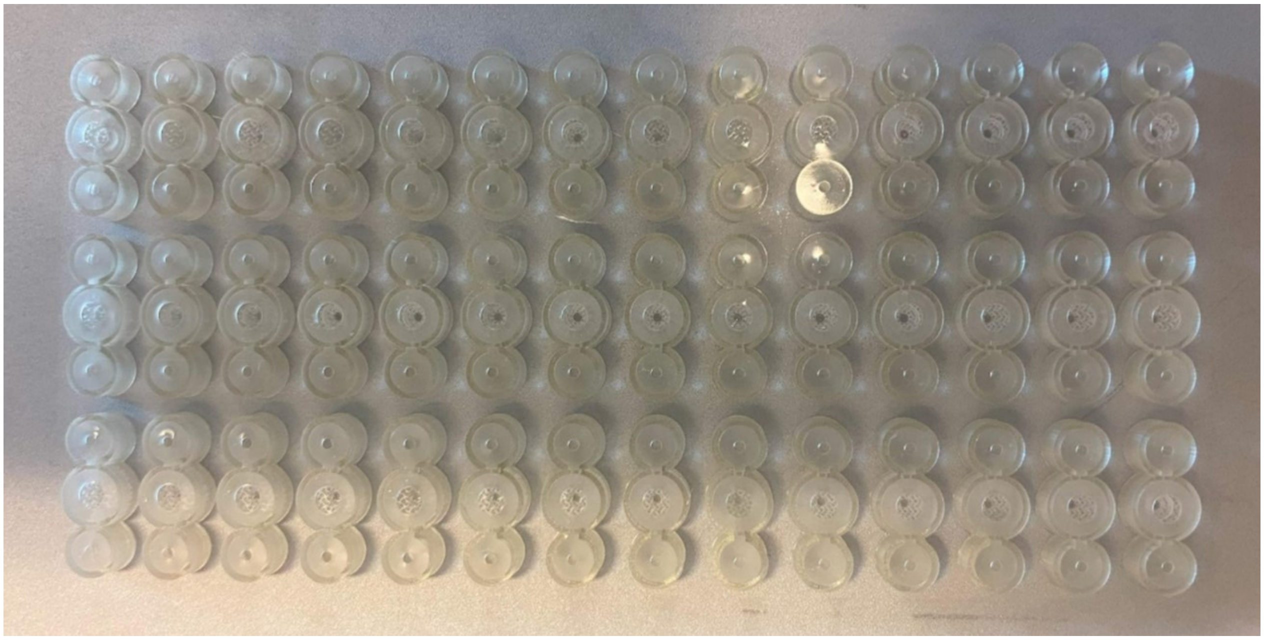

**Figure S15. 3D printing throughput for the OoC system**. Our process allowed for the fabrication of 42 chips in less than an hour, demonstrating significant production capabilities.

**Figure S16**

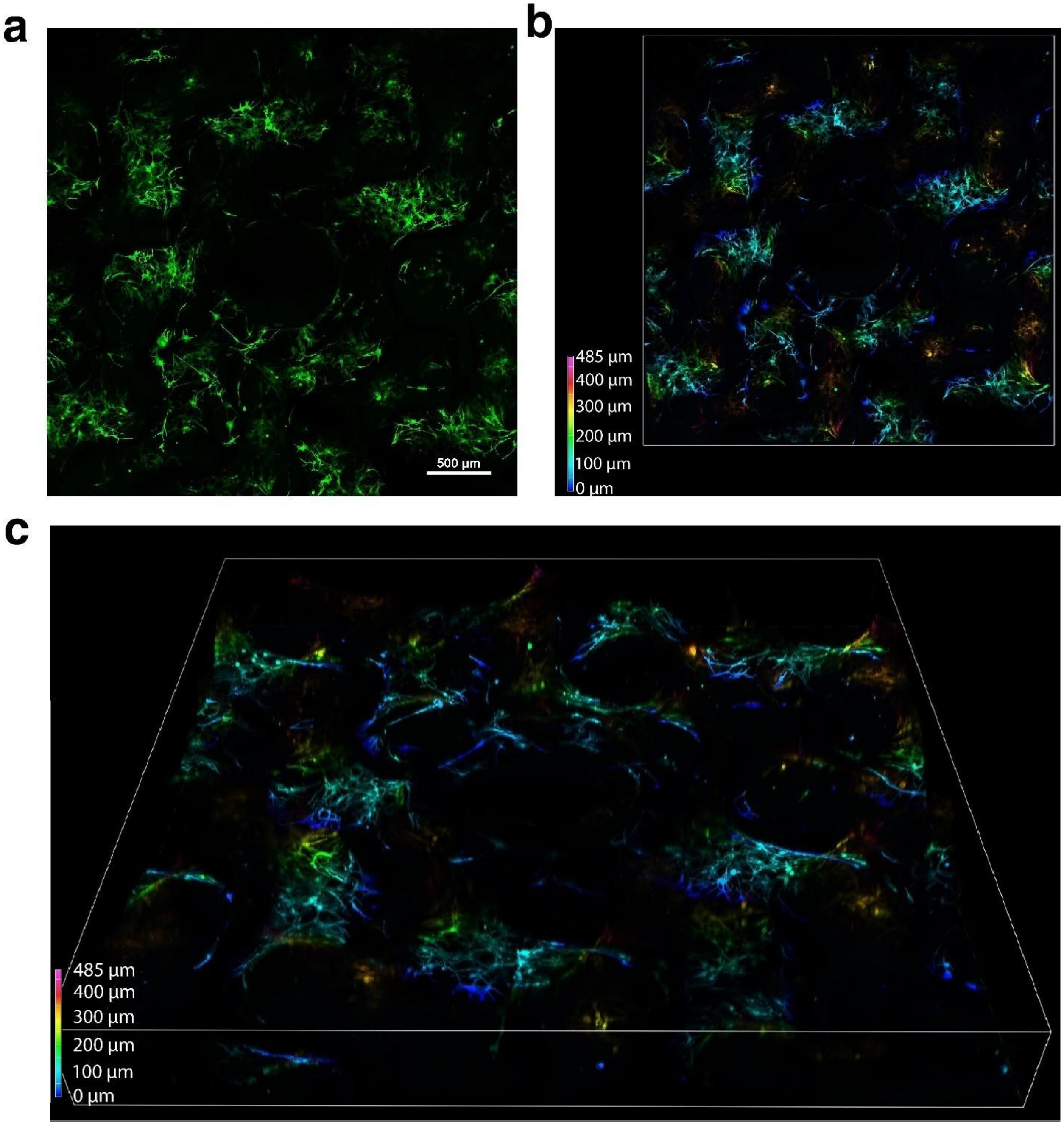


**Figure S16. Cultured IMR-90 cells within the 3D-printed gyroid structure. (a)** Fluorescence images of IMR-90 cells cultured within the gyroid, stopping at the CSV. **(b)** A 2D depth color map illustrating a height range of 480 µm. **(c)** A 3D depth color map representing the same height range of 480 µm.

Figure S17

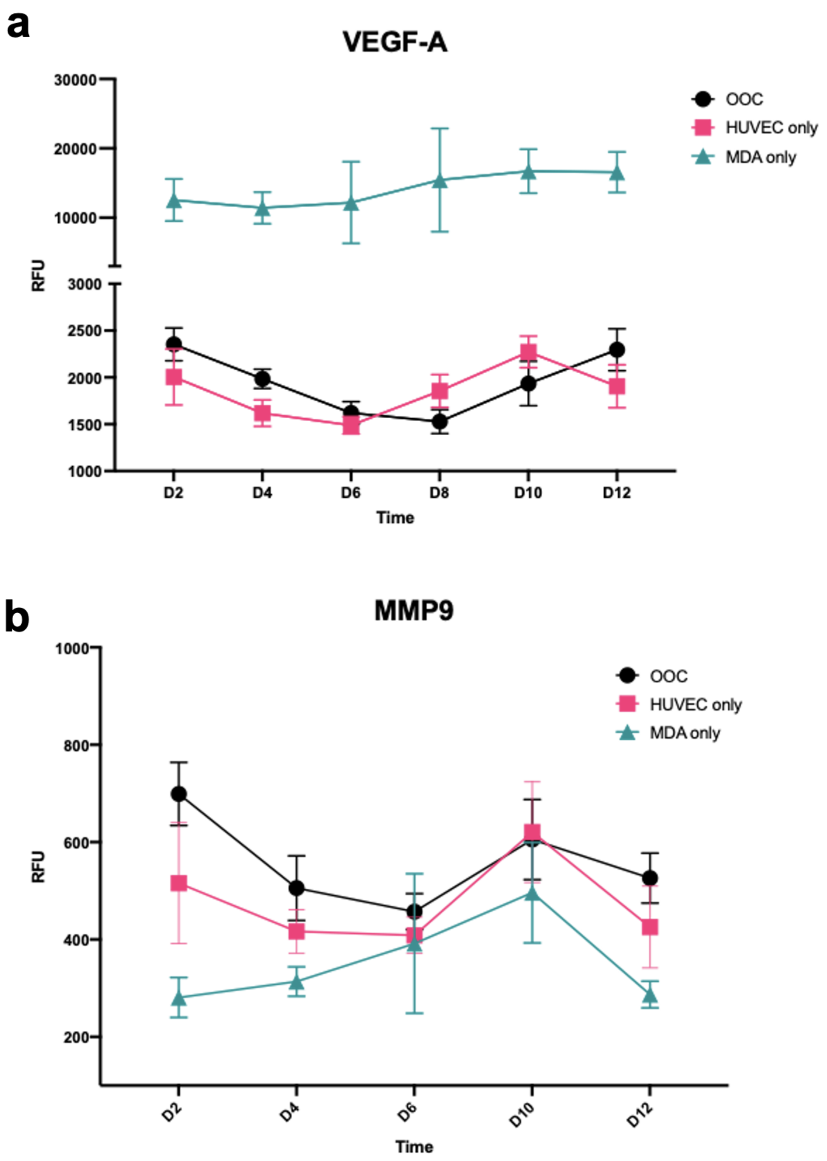


**Figure S17. Proteomic profile of VEGF-A and MMP9 in the OoC device.** Time course study of **(a)** VEGF-A and **(b)** MMP9 secretion from the OoC co-culture and mono-culture of cells, showing a consumption of VEGF-A and a rise in MMP9 in the OoC co-culture device. Data shows 3 biological repeats and 20 technical repeats for each condition and time course.
